# Targeting the host factor HGS-viral membrane protein interaction in coronavirus infection

**DOI:** 10.1101/2025.10.17.683077

**Authors:** Xubing Long, Rongrong Chen, Rong Bai, Buyun Tian, Yu Cao, Kangying Chen, Fuyu Li, Yiliang Wang, Yongjie Tang, Qi Yang, Liping Ma, Fan Wang, Maoge Zhou, Xianjie Qiu, Yongzhi Lu, Jie Zheng, Peng Zhou, Xinwen Chen, Qian Liu, Xuepeng Wei, Yongxia Shi, Yanhong Xue, Jincun Zhao, Wei Ji, Liqiao Hu, Jinsai Shang, Tao Xu, Zonghong Li

## Abstract

While current antivirals primarily target viral proteins, host-directed strategies remain underexplored. Here, we performed a genome-wide CRISPRi screening to identify the host protein, Hepatocyte Growth Factor-Regulated Tyrosine Kinase Substrate (HGS), as essential for the pan-coronaviruses infection both *in vitro* and *in vivo*. Mechanistically, HGS directly interacts with the viral membrane (M) protein, facilitating its trafficking to the ER-Golgi intermediate compartment (ERGIC) for virion assembly. Conversely, HGS deficiency caused M retention in the ER, blocking assembly. Leveraging this interaction, we designed M-derived peptides and screened over 5,000 FDA-approved drugs, identifying riboflavin tetrabutyrate (RTB). Both the peptides and RTB bind HGS and disrupt its interaction with the M protein, leading to M retention in the ER and subsequent blockade of virion assembly. These agents demonstrated broad anti-pan-coronavirus activity *in vitro* and *in vivo*. Collectively, our findings establish HGS as a druggable host target and identify RTB as a promising broad-spectrum antiviral candidate.

## Introduction

Host Targeting Agents (HTAs) have the potential to combat a broad spectrum of viruses, including newly emerging strains, by targeting host proteins essential for viral replication. This approach presents a higher genetic barrier to resistance compared to Viral Targeting Agents (VTAs), which directly target viral proteins, as host factors are not under the virus’s genetic control. HTAs can also simultaneously target multiple viruses, making them effective against co-infections. Notable examples of HTAs include Maraviroc, an FDA-approved drug that targets the CCR5 receptor to treat HIV-1 (1), Camostat, an inhibitor of TMPRSS2 and a clinically approved drug for chronic pancreatitis, can block the entry of SARS-CoV-2 (2), and Teicoplanin, a clinical antibiotic, also inhibited coronavirus entry by blocking cathepsin L (3). However, challenges arise, such as potential cytotoxicity and side effects associated with targeting host factors, as well as the possibility of viruses developing resistance by adapting their interactions with cellular factors. Additionally, the complexity of host-virus interactions necessitates advanced tools for screening and prediction.

Genome-wide CRISPR-Cas9 screening is a powerful tool for identifying critical host factors that regulate virus infection. Most studies have utilized cytopathic effect-based genome-wide CRISPR-Cas9 screening to identify key host factors involved in virus cell cycles (4–8). In the context of infection, insights are still mostly confined to early steps in the viral cycle, predominantly viral entry and replication. However, the identification of host factors involved in regulating virus assembly and egress through cytopathic effect-based screens has been limited. Targeting viral assembly and release provides another major antiviral strategy, which can synergize with strategies that suppress viral genome replication, leading to more effective overall antiviral activity. Here, using coronavirus as a model system, we established a genome-wide CRISPRi screening based on cell surface lysosome-associated membrane glycoprotein 1 (LAMP1) and identified a series of key host factors involved in virus assembly and egress. Subsequent research revealed HGS, a subunit of endosomal sorting complex required for transport (ESCRT), plays a vital role in pan-coronavirus infection *in vitro* and *in vivo*. HGS directly interacts with viral structural membrane (M) protein, facilitating its trafficking to the ERGIC for virion assembly. Importantly, the discovery of high-affinity peptides and an FDA-approved drug, RTB, targeting HGS have shown anti-pan-coronavirus activity *in vitro*, in ALI-cultured HBEs and *in vivo*, indicating the potential therapeutic significance of targeting HGS in combating coronavirus infections.

## Results

### Genome-wide CRISPRi screens identify HGS as a new host target for pan-coronavirus therapy

Numerous studies have suggested that coronavirus egress occurs through the lysosome, resulting in the upregulation of the lysosomal marker LAMP1 on the cell surface membrane (9–12). This observation led us to hypothesize that surface LAMP1 could potentially serve as a reliable indicator of coronavirus egress. To validate this hypothesis, we utilized specific anti-LAMP1 antibodies conjugated with APC and FITC to distinguish between cell surface and total LAMP1, where total LAMP1 serve as a control. Following infection with MHV, the ratio of cell surface LAMP1 to total LAMP1 was significantly elevated (Fig. S1A), consistent with previous findings (9, 10). Knockdown of *Arl8b*, a known critical host factor involved in the regulation of virion egress (10), led to a reduction in the cell surface LAMP1 to total LAMP1 ratio (Fig. S1A and S1B). This outcome indicates that the ratio of cell surface LAMP1 to total LAMP1 represents a suitable indicator of coronavirus egress.

Having established a suitable cellular model and fluorescence activated cell sorting (FACS)-based assay for indicator of coronavirus egress, we performed a genome-wide CRISPRi screen to identify the key host factors for coronavirus infection (Fig. 1A). A total of 434 candidate genes were substantially enriched and classified, which are related to vesicle-mediated transport and cytoskeleton (Fig. 1B and Table S1). Further validation of the top 24 enriched genes revealed that knockdown of most candidate genes led to a notable decrease in extracellular viral gRNA, while having no effect or, in some cases, even increasing intracellular viral gRNA levels at 16 h post-infection (hpi) (Fig. S1C and S1D). This pattern suggests that these candidate genes influence coronavirus assembly and egress, rather than replication, thereby affirming the specificity of the assay. Meanwhile, silencing the subunit of ESCRT, HGS, has significantly reduced coronavirus assembly and egress. Nevertheless, little is known about the role of HGS in coronavirus assembly and egress.

**Figure 1.**
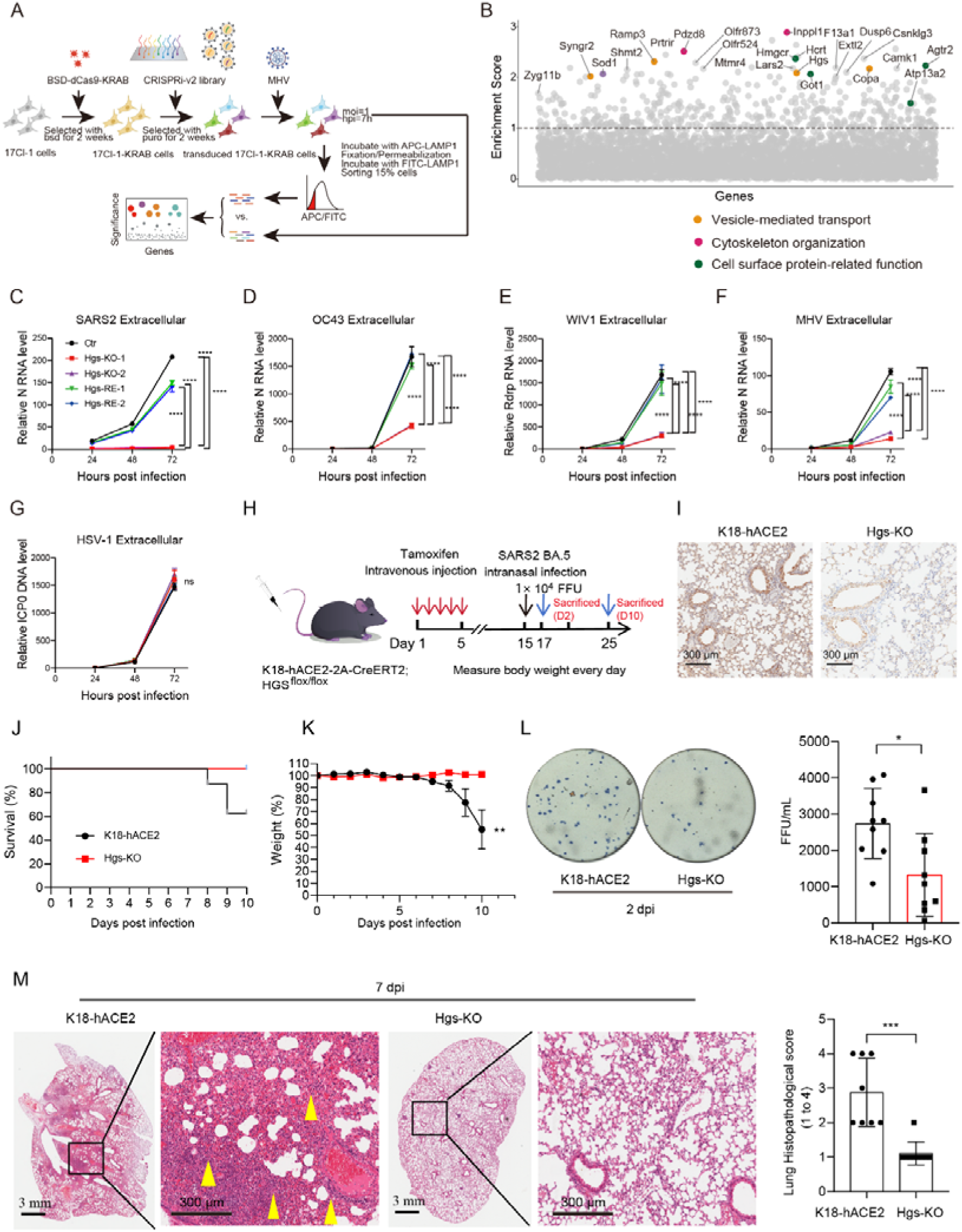
Genome-wide CRISPRi screens identify HGS as a new host target for pan-coronavirus therapy. (A) Schematic illustrating genome-wide CRISPRi screens based on surface membrane LAMP1 for the identification of host factors for coronavirus assembly and egress. dCas9-KRAB expressing 17Cl-1 cells were transduced with a genome-wide sgRNA library, selected with puromycin for two weeks, infected with MHV virus for 7 h (MOI = 1). The 15% cells with low ratio of cell surface LAMP1/total LAMP1 were enriched and sgRNA abundance was determined by next-generation sequencing. (B) Gene enrichment for CRISPRi screen of MHV infection. Enrichment scores were determined by MaGECK analysis and genes were colored by biological function. Dotted line indicates log10(Enrichment Score) = 1. All genes and their enrichment scores can be found in Table S1. (C-G) RT-qPCR analysis of extracellular SARS-CoV-2 (C), HCoV-OC43 (D), WIV1 (E), MHV (F) viral gRNA and HSV-1 (G) viral gDNA levels in the Ctr, two *HGS*-KO clones and their respective *HGS*-rescued cells (hpi = 24 h, 48 h, 96 h respectively, MOI = 1). N = 3 independent biological replications. (H) Schematic illustrating SARS-CoV-2 infection in the mice of conditional *Hgs* KO specifically in hACE2-expressing cells. K18-hACE2-T2A-CreERT2/HGS-floxed mice were intraperitoneal injection of tamoxifen to induce Cre-mediated recombination for 5 days. After 15 days of tamoxifen treatment, the mice were intranasally infected with SARS-CoV-2 Omicron BA.5 (1 × 10^4^ focus forming unit, FFU) for 10 days. (I) Immunohistochemical staining of *Hgs* expression in the lung of K18-hACE2-T2A-CreET2/HGS-floxed mice treated with (Hgs-KO) or without (K18-hACE2) tamoxifen. Scale bar[=[300[μm. (J-K) Survival curves (J) and body weight changes (K) were analyzed after infection with SARS-CoV-2 Omicron BA.5 (N = 9 for K18-hACE2 group and N = 8 for Hgs-KO group). (L) Viral titration by focus forming assay (FFA) with the supernatant of homogenized lung tissues on day 2 (N = 9 for each group). (M) Histopathology of formalin-fixed and hematoxylin and eosin (HE)-stained lung tissues on day 7. Quantitative analysis of pathological severity scores based on the percentage of affected area in lung tissues (N = 9 for K18-hACE2 group and N = 8 for Hgs-KO group) Data are the mean ± SD. Significance testing for (C-G) was performed with 1-way ANOVA and Tukey’s multiple comparison test. Significance testing for (K-M) was performed with an unpaired t test. *P ≤ 0.05, **P ≤ 0.005, ***P ≤ 0.0005, ****P ≤ 0.0001, ns, no significance.

To investigate the role of HGS in the coronavirus infection, *HGS* knockout (KO) Huh7.5.1 and 17Cl-1 cell lines generated using CRISPR-Cas9 were subjected to various coronaviruses infection, including SARS-CoV-2, HCoV-OC43, a bat coronavirus WIV1 and MHV. Quantitative real-time PCR (RT-qPCR) results showed that extracellular viral gRNA levels were dramatically decreased in *HGS*-KO cells but restored in *HGS*-rescued cells at 72 hpi (Fig. 1C-1F). Notably, extracellular viral gDNA levels were unchanged in *HGS*-KO and rescued cells following infection with the DNA virus HSV-1 (Fig. 1G). These results indicate that HGS functions as a critical host factor for pan-coronavirus infection.

We further investigated the role of HGS in coronavirus infection *in vivo*. Due to the embryonic lethality of global *Hgs* KO mice(13), we employed a conditional gene-targeting approach to generate mice with selective HGS deficiency in hACE2-expressing cells (Fig. 1H and 1I). Strikingly, HGS deficiency markedly attenuated body weight loss and significantly enhanced the survival rate of SARS-CoV-2 Omicron BA.5-infected mice (Fig. 1J and 1K). Furthermore, HGS deficiency reduced viral titers in the lungs on day 2 post-infection (Fig. 1L). Histopathological analysis further revealed that lung inflammation was alleviated in HGS-deficient mice (Fig. 1M). These results indicate that HGS functions as a critical host factor for SARS-CoV-2 infection *in vivo*.

In addition, we selectively inactivated HGS in the liver by intravenously injecting liver-tropic adeno-associated virus (AAV) containing sgRNA specifically targeting the *Hgs* gene into spCas9 knock-in mice (Fig. S2A). Immunoblotting (IB) results revealed a significant decrease in *Hgs* expression in the liver, while no significant change was observed in the lung, as compared to control mice treated with scrambled sgRNA (Fig. S2B and S2C), which confirmed the tissue-specific tropism of the AAV. Subsequently, the mice were intranasally infected with MHV for 5 days. Body weight loss did not show significant differences in the liver-specific *Hgs* silencing mice compared to control mice (Fig. S2D). However, viral gRNA levels and virus titers were significantly decreased in the liver but not the lung of the liver-specific *Hgs* silencing mice (Fig. S2E-S2G). Histological analysis further indicated that liver damage was alleviated in the liver-specific *Hgs* silencing mice, while acute tissue damage in the lung was not significantly improved (Fig. S2H and S2I). These results collectively demonstrate the essential role of HGS in MHV infection *in vivo*.

### HGS controls pan-coronavirus assembly

We further investigate whether HGS plays a role in the virus genome replication or virion assembly and egress. HGS deficiency resulted in a significant decrease in secreted viral gRNA levels, while intracellular viral gRNA levels remained unaffected after various coronaviruses infection for 16 h, including MHV, SARS-CoV-2, HCoV-229E, HCoV-OC43, HCoV-NL63 and a bat coronavirus WIV1 (Fig. S3). Conversely, the rescued expression of *HGS* in the *HGS-*KO cells restored secreted viral gRNA levels without altering intracellular viral gRNA levels (Fig. S3). Furthermore, the intracellular protein level of viral structural protein N was unchanged in both *HGS*-KO and rescued cells (Fig. S3). These findings strongly indicate that HGS plays a regulatory role in virion assembly and egress, rather than genome replication.

To investigate the role of HGS in regulating coronavirus assembly or release, intracellular and extracellular virus titers were assessed using a virus plaque assay. The results showed a significant reduction in both intracellular and extracellular virus titers in *Hgs*-KO 17Cl-1 cells infected with MHV, and an increase in both intracellular and extracellular virus titers in the *Hgs*-rescued cells (Fig. 2A and 2B), suggesting that HGS predominantly influences coronavirus assembly. Consistently, *HGS* deficiency also decreased other coronavirus assembly, including SARS-CoV-2, HCoV-229E, HCoV-OC43, HCoV-NL63 and WIV1 (Fig. 2C-2G). Furthermore, transmission electron microscope (TEM) observations revealed intracellular mature virions were significantly decreased in *Hgs*-KO cells compared with control cells, while DMVs, the replication organelles, were not significantly changed (Fig. 2H-2K), suggesting that the absence of HGS does not affect replication but impedes virion assembly.

**Figure 2.**
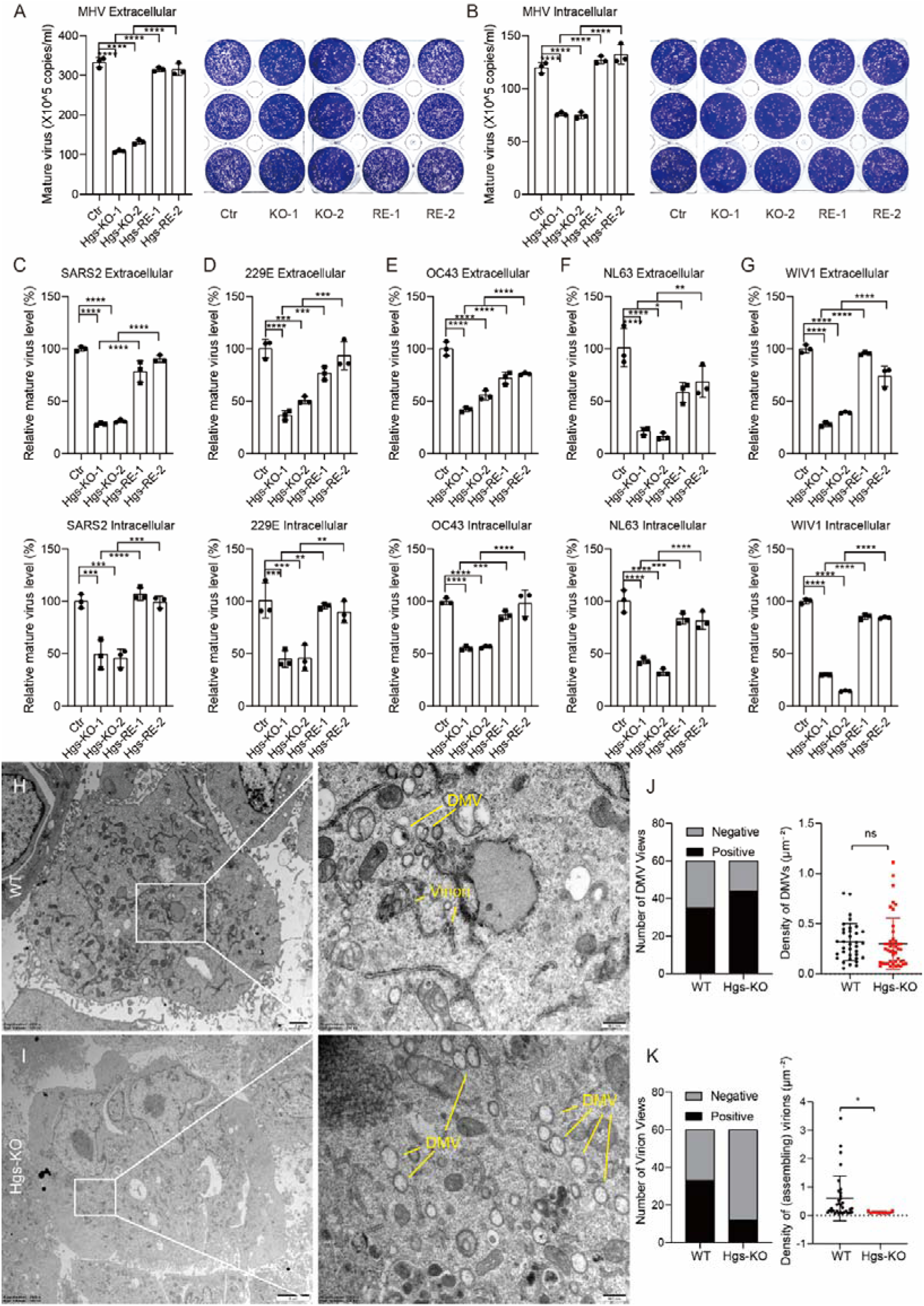
HGS is essential for virion assembly. (A-B) Plaque assay analysis of extracellular (A) and intracellular (B) mature MHV titer levels in the Ctr, two *Hgs*-KO clones and their respective *Hgs*-recused 17Cl-1 cells (hpi = 16 h, MOI = 1). N = 3 independent biological replications. (C-G) RT-qPCR analysis of extracellular (upper panel) and intracellular (lower panel) mature virion of SARS-CoV-2 (C), HCoV-OC43 (D), HCoV-229E (E), HCoV-NL63 (F) and WIV1 (G) in the Ctr, two *HGS*-KO clones and their respective *HGS*-recused Huh7.5.1 cells (hpi = 24 h, MOI = 1). The extracellular and intracellular mature virion from Ctr, two *HGS*-KO clones and their respective *HGS*-recused Huh7.5.1 cells were subjected to infect WT Huh7.5.1 cells for 6 h, The intracellular viral gRNA levels were examined to indicate the mature virion level. N = 3 independent biological replications. (H-K) TEM analysis of MHV infected Ctr (H) and *Hgs*-KO (I) 17Cl-1 cells (hpi = 24 h, MOI = 1). (J) Quantitative analysis of the number of DMV-positive views (left panel) and the number of DMVs per unit cytoplasmic area (right panel). (K) Quantitative analysis of the number of virion-positive views (left panel) and the number of vesicle-contained virions per unit cytoplasmic area (right panel). Data are the mean ± SD. Significance testing for (A-G) was performed with 1-way ANOVA and Tukey’s multiple comparison test. Significance testing for (J-K) was performed with an unpaired t test. *P ≤ 0.05, **P ≤ 0.005, ***P ≤ 0.0005, ****P ≤ 0.0001, ns, no significance.

In addition, various coronavirus VLPs containing M/E/N proteins were utilized to investigate coronavirus assembly, including SARS-CoV-2, MERS, HCoV-OC43 and HCoV-HKU1. Overexpression of *HGS* significantly increased the ratio of secreted to intracellular N protein (Fig. S4A-S4D). In contrast, HGS deficiency markedly reduced this ratio, while HGS rescue restored it to levels comparable to those observed in control cells (Fig. S4E-S4H). All these results strongly support the role of HGS in regulating pan-coronavirus assembly.

### HGS directly interacts with M protein and facilitates its trafficking to ERGIC for virion assembly

We went on to investigate the mechanism by which HGS regulates virion assembly. Previous studies have established that HGS interacts with multiple cellular proteins and modulates their intracellular trafficking and sorting (14, 15). Given that the coronavirus M protein serves as the central organizer of virion assembly, with its proper subcellular localization being essential for this process (16–18), we hypothesized that HGS might regulate M protein trafficking to facilitate viral particle formation. In wild-type (WT) cells, M protein was co-localized with an ERGIC marker, consistent with previous studies demonstrating that coronavirus assembly occurs in the ERGIC (19, 20). However, in *HGS*-KO cells, the M protein co-localized with an ER marker, but not with an ERGIC marker (Fig. 3A). Consistently, similar results were observed in SARS-CoV-2-infected cells (Fig. 3B), indicating that HGS facilitates M protein trafficking to ERGIC for virion assembly.

**Figure 3.**
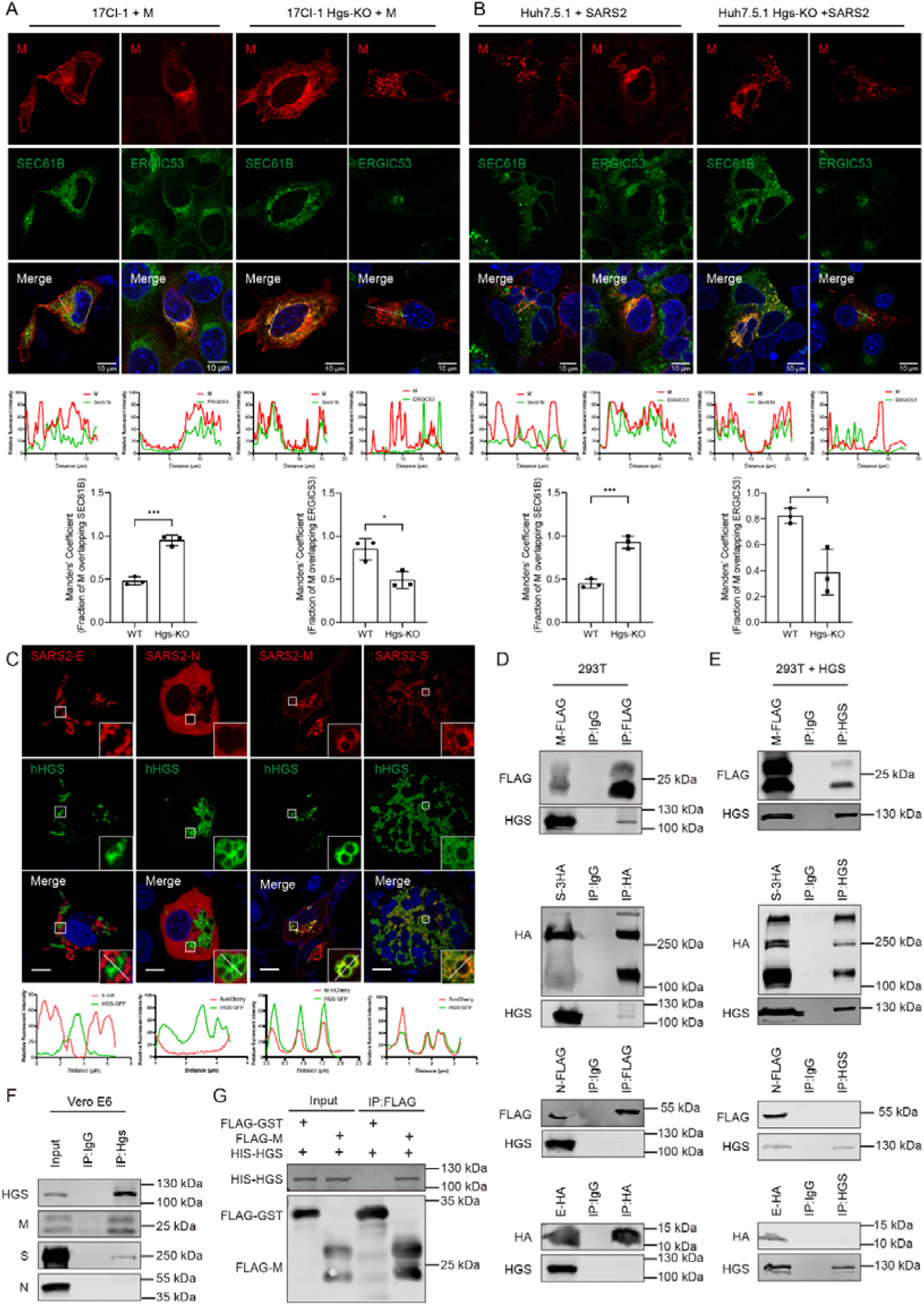
HGS directly interacts with M protein and facilitates its trafficking to ERGIC. (A) Representative IF analysis of the colocalization of SARS-CoV-2 M with SEC61B or ERGIC53 in WT and *Hgs*-KO 17Cl-1 cells. Trace outline is used for line-scan analysis of the relative fluorescence intensity of SARS-CoV-2 M with SEC61B and ERGIC53. Scale bar, 10 μm. Quantitative image analysis of M-SEC61B/ERGIC53 colocalization using Manders’ Coefficient. N = 3 independent biological replications. (B) Representative IF analysis of the colocalization of SARS-CoV-2 M with SEC61B in SARS-CoV2 Omicron BA.5-infected Huh7.5.1 WT and *HGS*-KO cells (MOI = 0.5). Trace outline is used for line-scan analysis of the relative fluorescence intensity of SARS-CoV-2 M with SEC61B. Scale bar, 10 μm. Quantitative image analysis of M-SEC61B/ERGIC53 colocalization using Manders’ Coefficient. N = 3 independent biological replications. (C) Representative IF analysis of the colocalization of HGS with SARS-CoV-2 M, S, E, and N in Vero E6 cells. Trace outline is used for line-scan analysis of the relative fluorescence intensity of HGS with SARS-CoV-2 M, S, E, and N. Scale bar, 10 μm. N = 3 independent biological replications. (D) Co-IP analysis of the interaction of SARS-CoV-2 M, S, E, N with HGS. SARS-CoV-2 M, S, E and N Proteins were individually transiently expressed in HEK293T cells. Immunoprecipitates pulled down by FLAG or HA antibody were analyzed by IB with the indicated antibodies. IgG IP was used as a negative control. Input represents 5% of the total cell extract used for immunoprecipitation. Molecular weights are in kDa. N = 3 independent biological replications. (E) Co-IP analysis of the interaction of HGS with SARS-CoV-2 M, S, E, and N. Proteins were transiently expressed in HEK293T cells, and immunoprecipitates pulled down by HGS antibody were analyzed by IB with the indicated antibodies. IgG IP was used as a negative control. Input represents 5% of the total cell extract used for immunoprecipitation. Molecular weights are in kDa. N = 3 independent biological replications. (F) Co-IP analysis of the interaction of HGS with M, S and N in the SARS-CoV-2 infected Vero-E6 cells. Vero-E6 cells infected SARS-CoV-2 for 24 h were lysed, and immunoprecipitates pulled down by HGS antibody were analyzed by IB with the indicated antibodies. IgG IP was used as a negative control. Input represents 5% of the total cell extract used for immunoprecipitation. Molecular weights are in kDa. N = 3 independent biological replications. (G) *In vitro* pull-down analysis of the interaction between HGS and SARS-CoV-2 M. Purified HGS and SARS-CoV-2 M proteins were subjected to pull-down assay, and pull-down samples by FLAG antibody were analyzed by IB with the indicated antibodies. FLAG-GST protein was used as a negative control. Input represents 5% of the total proteins used for pull-down. Molecular weights are in kDa. N = 3 independent biological replications. Data are the mean ± SD. Significance testing for (A-B) was performed with an unpaired t test. *P ≤ 0.05, **P ≤ 0.005, ***P ≤ 0.0005, ****P ≤ 0.0001, ns, no significance.

We further explore how HGS facilitates M protein trafficking to ERGIC. Co-expressing of HGS with individual SARS-CoV-2 viral structural protein M, E, N and S showed that HGS co-localized with viral structural protein M and S, but not with E and N (Fig. 3C). Subsequent co-immunoprecipitation (IP) results showed that HGS was immunoprecipitated by viral structural protein M and S, not by E and N (Fig. 3D). Conversely, viral structural protein M and S, but not E and N were immunoprecipitated by HGS (Fig. 3E). These results showed that HGS interacted with viral structural protein M and S, which were further confirmed by endogenous HGS IP in SARS-CoV-2 infected cells (Fig. 3F). Moreover, purified HGS protein and M protein were subjected to pull down assay. IB results indicated direct interaction between M protein and HGS, as HGS protein was pulled down by M protein but not by the GST-control protein (Fig. 3G). Taken together, all these results showed that HGS directly interacts with viral structural protein M.

Subsequently, we investigated whether HGS could interact with other coronavirus M protein. Immunofluorescence (IF) analysis revealed that HGS co-localized with all coronavirus M protein, including those of MHV, HCoV-229E, HCoV-OC43, HCoV-NL63, HCoV-HKU1, MERS, SARS-CoV-1, and WIV1 (Fig. S5A).

Additionally, different HGS species were observed to co-localize with SARS-CoV-2 M protein (Fig. S5B). These findings collectively suggest that the interaction between HGS and M protein is conserved across diverse coronaviruses.

To identify the specific domain within HGS required for the interaction with M, various truncations of HGS were generated, including HGS (1-166), HGS (1-215), HGS (1-290), HGS (1-390) and HGS (1-509) (Fig. S6A). IF analysis showed that HGS (1-290), HGS (1-390) and HGS (1-509) were capable of forming vesicular structures and co-localized with M protein, similar to WT HGS (Fig. S6B). Consistently, co-IP results showed that HGS (1-290), HGS (1-390) and HGS (1-509), but not HGS (1-166) and HGS (1-215) were immunoprecipitated by M protein (Fig. S6C). These results demonstrated the essential role of HGS (1-290) region in mediating its interaction with M protein.

To identify the specific domain within M required for the interaction with HGS, various truncations of M protein were generated, including a transmembrane domain truncation (1-118) and an intravirion domain truncation (Δ19-100) (Fig. S6D). IF results showed that HGS co-localized with M intravirion domain truncation (Δ19-100), but not with M transmembrane domain truncation (1-118) (Fig. S6E). Furthermore, the purified HGS protein was pulled down by the M intravirion protein but not by the FLAG-control protein (Fig. S6F), indicating a direct interaction between HGS and the M intravirion domain. Subsequently, we sought to further narrow the specific binding domain within the M intravirion domain with HGS. The M intravirion domain comprises eight β-sheet domains, and we sequentially generated four truncations that deleted these domains: M (Δ118-134), M (Δ135-161), M (Δ162-189), and M (Δ190-204) (Fig. S6D). IF results revealed that only M (Δ135-161) did not co-localize with HGS (Fig. S6E), indicating the essential role of the M (135-161) domain in its interaction with HGS. Additionally, both M (135-161)-mCherry and M (135-146)-mCherry showed complete co-localization with HGS (Fig. S6E), further confirming a direct interaction between the M (135-146) domain and HGS. Collectively, these results show HGS directly interacts with M protein and facilitates its trafficking to ERGIC for virion assembly.

### M-derived peptides targeting HGS alleviate the coronavirus infection *in vitro*, in ALI-cultured HBEs and *in vivo*

The pioneering results promoted us to investigate whether the M binding domain could serve as a dominant-negative truncation to disrupt the interaction between HGS and M, thereby preventing coronavirus assembly. To verify this hypothesis, we observed a significant decrease in extracellular virus titer in SARS-CoV-2 M (135-161) and M (135-146) overexpressing 17Cl-1 cells infected with MHV compared to FLAG-control cells (Fig. S7A), indicating that M-derived peptides may impede the HGS-M interaction and hinder coronavirus infection. To expand on the clinical applicability, the M (135-161) and M (135-146) peptides were designed to fuse with HIV-TAT, a cell-penetrating peptide known to facilitate cellular uptake, which we called M161 and M146 here, respectively. Surface plasmon resonance (SPR) results showed that the binding affinity between M146 and M161 with HGS reached to 5.73 nM and 3.84 nM, respectively (Fig. S7B and S7C). The M146 and M161 peptides could significantly block the interaction between HGS and M protein (Fig. S7D). Cell viability assessments demonstrated that M146 and M161 exhibited no significant cytotoxicity up to 10^4^ nM (Fig. S7E-S7G). Strikingly, M146 and M161 treatment significantly decreased the extracellular viral gRNA levels but not affected the intracellular viral gRNA levels in a dose-dependent manner after infection with MHV for 24 h, however, no significant change of extracellular and intracellular viral gRNA level was observed in the control peptide (Fig. S7H and S7I). When coronavirus infection continues for 72 h, M146 and M161 treatment at 10^4^ nM strikingly decreased the MHV virus infection, but not affected the HSV-1 virus infection (Fig. S7J and S7K). Furthermore, when M146 and M161 were mutated to decrease their binding affinity to HGS, the inhibition of virus release was consistently reduced (Fig. S7L-S7P). Since HGS is a key host factor for pan-coronavirus assembly, we extended our analysis to various coronaviruses, including SARS-CoV-2, HCoV-NL63, HCoV-229E, HCoV-OC43 and WIV1. Treatment with M146 and M161 at 10^4^ nM for 24 h markedly decreased extracellular viral gRNA levels across all coronavirus-infected cells, while intracellular viral gRNA levels remained unchanged (Fig. S7Q-S7U), suggesting that targeting HGS could effectively mitigate coronaviruses assembly.

The airway ciliated epithelial cells are the main targets for initial coronaviruses infections, and spread of progeny to neighboring cells is vital for coronavirus infection (21). Therefore, we differentiated primary HBE cells in ALI cultures to form bronchial epithelial barrier containing ciliated, goblet, and basal cells (Fig. S8A and S8B). The fully differentiated ALI-cultured HBEs treated with M146 and M161 were inoculated with various coronaviruses, including SARS-CoV-2, HCoV-229E, HCoV-OC43 and HCoV-NL63. IF analysis showed that N proteins were most seen in ciliated cells (Fig. S8C), consistent with previous results (21). Strikingly, M146 and M161 treatment significantly decreased the N protein levels (Fig. S8D-S8G). Consistently, RT-qPCR analysis of mucus layer revealed that the viral gRNA was dramatically decreased after treatment with M146 and M161, compared to the FLAG-control (Fig. S8H-S8K). Taken together, targeting HGS could effectively mitigate coronavirus infections in ALI-cultured HBEs.

To determine the function of HGS-targeted peptides *in vivo*, peptide M146 and M161 were modified with d-retroinverso (DRI) isoform to enhance the peptide potency. Several DRI-modified peptides have reported to be well-tolerated and therapeutically effective in clinical trials (22–24). The mice pre-injected with M146-DRI and M161-DRI were infected with MHV. Tissue distribution analysis showed that the M146-DRI peptide mainly localized in the lung after intravenous injection (Fig. S8M). RT-qPCR and virus plaque analysis showed that the viral gRNA and virus titer in the lung was significantly decreased after treatment with M146-DRI and M161-DRI (Fig. S8L-S8P). Consistently, histological analysis indicated that the lung injure was alleviated in the mice treatment with M146-DRI and M161-DRI (Fig. S8Q). Together, these results indicate that targeting HGS could alleviate the coronavirus infection *in vivo*.

### Small-molecule screening reveals RTB disrupts HGS-M interaction

To identify the small molecules that disrupt the interaction between of HGS and M protein, the work flow of compounds screening based on fluorescence polarization (FP) assay was applicated in a preliminary screening of an in-house drug library of over 5000 compounds with known structure (Fig. 4A and 4B). By selecting compounds exhibiting more than 60% inhibition, a hit list of five candidates (Eltrombopag, RTB, Citropten, Ethacridine lactate and 5-amino-2-methoxypyridine) was identified (Fig. 4C). We further measured the IC_50_ values of the hit compounds identified from high-throughput screening, among which RTB exhibiting the most potent inhibitory activity (IC_50_ = 0.58 μM) (Fig. 4D). Cell viability assays confirmed that all five compounds exhibited minimal cytotoxicity (CC_50_ >20 μM). Among these, RTB most effectively rescued coronavirus-induced cytopathic effects, with an EC_50_ of 3.48 μM. This antiviral potency was benchmarked against remdesivir (RDV, EC_50_ = 0.69 μM) (Fig. S9). This pilot screening further illustrated that the FP-based assay can be used as a robust HTS assay for the discovery of HGS protein inhibitors.

**Figure 4.**
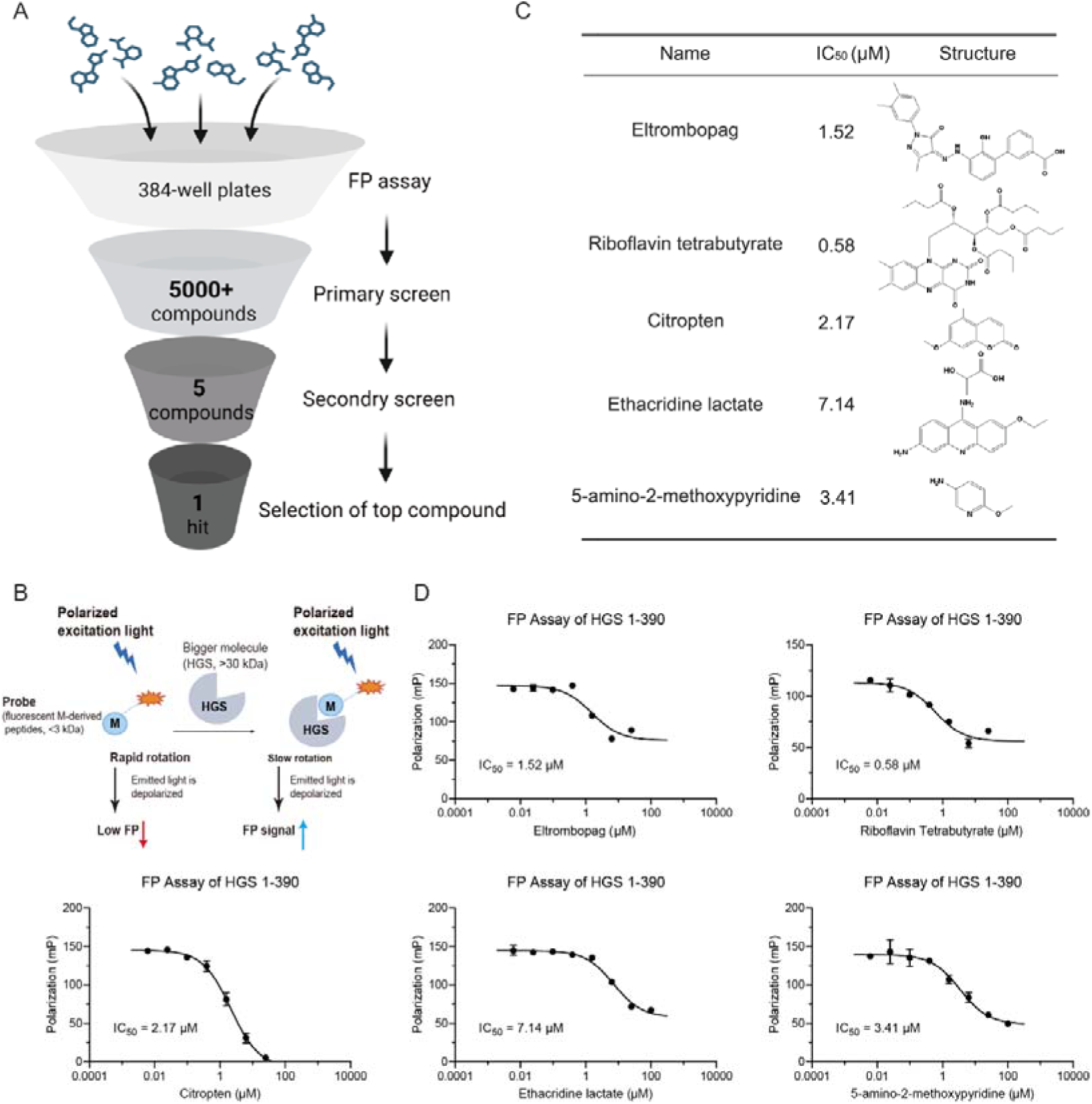
Discovery of small-molecule inhibitor targeting HGS. (A) The flowchart of FP-based HTS assay. (B) Schematic representation of the FP-based HTS assay. (C) Half of inhibition concentration (IC_50_) and structure of 5 compounds, which showed the best inhibitory activity among all the compounds. (D) The IC_50_ values of the top 5 hits were shown in the inhibition curves of HGS (1-390). Data are the mean ± SD and analyzed in GraphPad Prism 9.3.

### RTB exhibits anti-coronavirus activity *in vitro*, in ALI-cultured HBEs and *in vivo*

We further explore the role of RTB in the various coronaviruses infection. RTB demonstrated broad-spectrum anti-coronaviral infection with the following EC_50_ values, 14.85 μM for SARS-CoV-2 WT, 4.09 μM for SARS-CoV-2 Omicron BA.5, 8.46 μM for HCoV-NL63, 11.22 μM for MHV, 13.88 μM for HCoV-OC43, 6.11 μM for HCoV-229E, and 3.53 μM for WIV1, respectively (Fig. 5A-5G). Notably, all EC_50_ values for RTB were higher than those of RDV. In fully differentiated ALI cultures of HBEs, RTB produced significant, dose-dependent inhibition of HCoV-OC43, HCoV-229E, HCoV-NL63, and WIV1 infection (Fig. 5H-5K).

**Figure 5.**
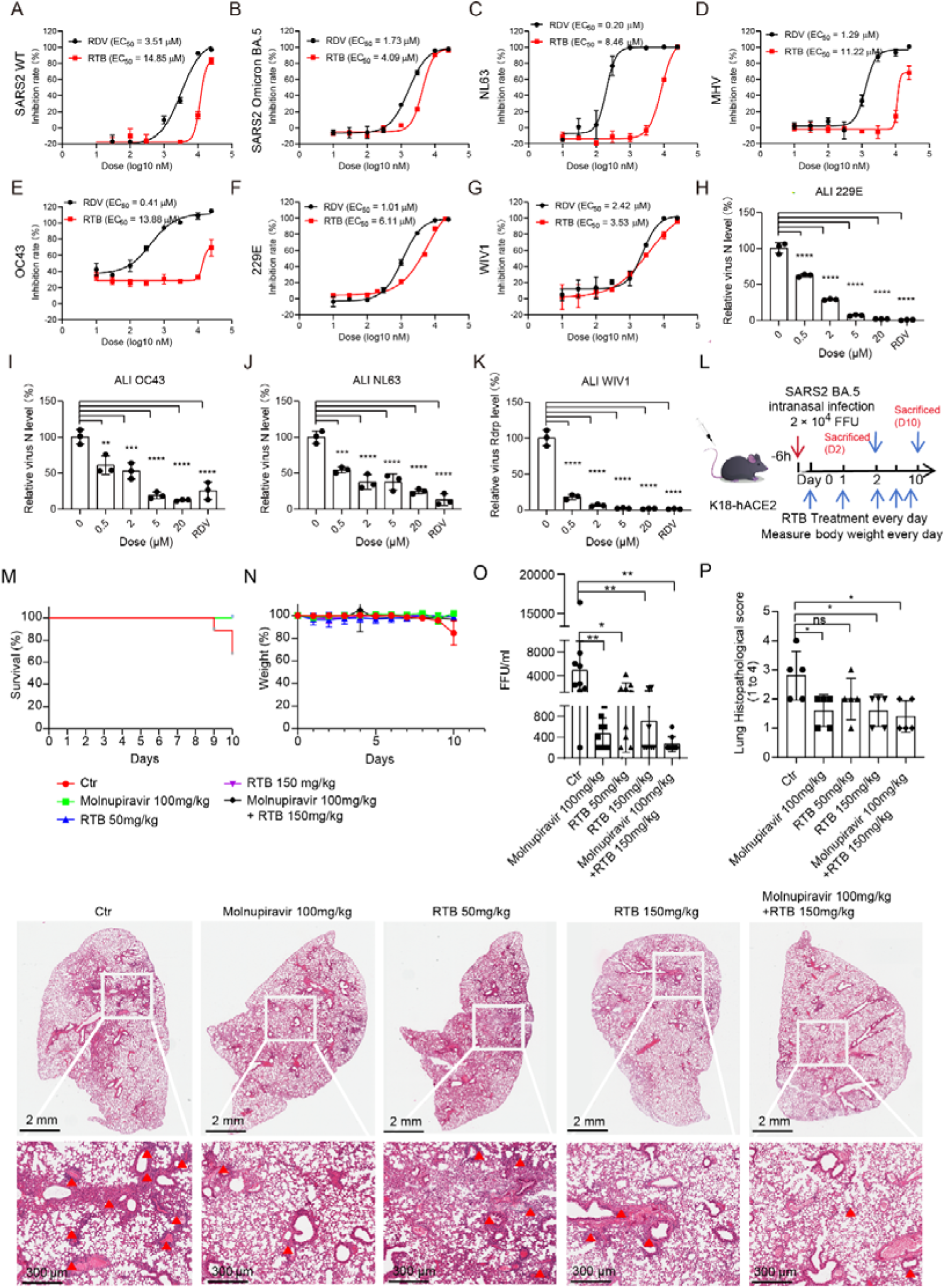
RTB exhibits anti-coronavirus activity *in vitro*, in ALI-cultured HBEs and *in vivo*. (A-G) Dose titration of RTB and RDV for anti-coronavirus activity. Various concentrations of the RTB and RDV were used to assess their ability to inhibit various coronaviruses infection by indirect immunofluorescence assay (IFA). EC_50_ value of antiviral ability are indicated. (A) SARS-CoV-2 wild-type (MOI = 0.1, hpi = 24 h), (B) SARS-CoV-2 Omicron BA.5 (MOI = 0.1, hpi =24 h), (C) HCoV-NL63 (MOI = 0.1, hpi = 48 h), (D) MHV (MOI = 0.1, hpi = 24 h) , (E) HCoV-OC43 (MOI = 0.1, hpi = 48 h), (F) HCoV-229E (MOI = 0.1, hpi = 48 h), (G) WIV1 (MOI = 0.1, hpi = 48 h). Inhibition Rate = [1- (Infection Rate of the Test Compound - Cell Control) / Infection Rate of the Virus Control] x 100%. Following the analysis of inhibition rates, the EC_50_ is determined using a four-parameter fitting process. (H-K) RTB alleviates the coronavirus infection in ALI-cultured HBEs. RT-qPCR analysis of extracellular viral gRNA levels in the HCoV-229E (MOI = 1) (H), HCoV-OC43 (MOI = 1) (I), HCoV-NL63 (MOI = 1) (J), WIV1 (MOI = 1) (K) infected ALI-cultured HBEs treatment with different doses of RTB or 5 μM RDV for 96 h. (L) Schematic illustrating SARS-CoV-2 Omicron BA.5 infection in the K18-hACE2 mice treatment with or without RTB. SARS-CoV-2 Omicron BA.5-infected mice (2 x 10^4^ FFU) were treated with PBS, Molnupiravir (100 mg/kg body weight), RTB (50 or 150 mg/kg body weight), and combining Molnupiravir (100 mg/kg body weight) with RTB (150 mg/kg body weight) every 24 h for 2 or 10 days. (M-N) Survival curves (M) and body weight changes (N) were analyzed after infection with SARS-CoV-2 Omicron BA.5 (N = 9 for each group). (O) Viral titration by FFA with the supernatant of homogenized lung tissues on day 2 (N = 9 for each group) (P) Histopathology of formalin-fixed and HE-stained lung tissues on day 2. Quantitative analysis of pathological severity scores based on the percentage of affected area in lung tissues (N = 5 for each group) Data are the mean ± SD. Significance testing for (H-K, O) was performed with 1-way ANOVA and Tukey’s multiple comparison test. *P ≤ 0.05, **P ≤ 0.005, ***P ≤ 0.0005, ****P ≤ 0.0001, ns, no significance.

To determine the function of RTB *in vivo,* K18-hACE2 C57BL/6 mice were infected with the SARS-CoV-2 Omicron BA.5 variant with RTB administered at the onset of infection (Fig. 3L). RTB treatment significantly improved survival and markedly reduced body weight loss in infected mice (Fig. 5M and 5N). Consistently, RTB substantially lowered pulmonary viral titers and alleviated lung inflammation (Fig. 5O and 5P).

To further explore whether RTB could be an effective antiviral against other coronaviruses *in vivo*, MHV infected WT C57BL/6 mice and 229E infected K18-hACE2 C57BL/6 mice were administered RTB at the onset of infection (Fig. S10). Remarkably, RTB treatment significantly reduced viral gRNA and lowered pulmonary viral titers in both models. Critically, RTB markedly attenuated virus-induced lung injury (Fig. S10). Collectively, these results suggest that RTB could alleviate various coronaviruses infection *in vivo*.

### RTB directly targets HGS

To investigate the conformational change of HGS induced by RTB, we performed hydrogen-deuterium exchange mass spectrometry (HDX-MS), a widely established technique for probing protein structure, dynamics, folding, and interactions. The different deuterium uptake rates of HGS (1-390) upon RTB binding revealed the specific regions that interact with RTB as well as conformational changes induced by RTB binding. Two regions showed significantly different deuterium incorporation in treated HGS with or without RTB. Interestingly, one region, covering residues 167-175, showed an average 6% increase in deuterium uptake, indicating that the binding of RTB to HGS may have altered the conformational change in this region making it more susceptible to exchange with deuterium water. The other region (residues 209-216) exhibited an approximate 10% decrease in deuterium uptake upon HGS binding to RTB, suggesting that this region may be affected by the binding between RTB and HGS (Fig. 6Α-6D).

**Figure 6.**
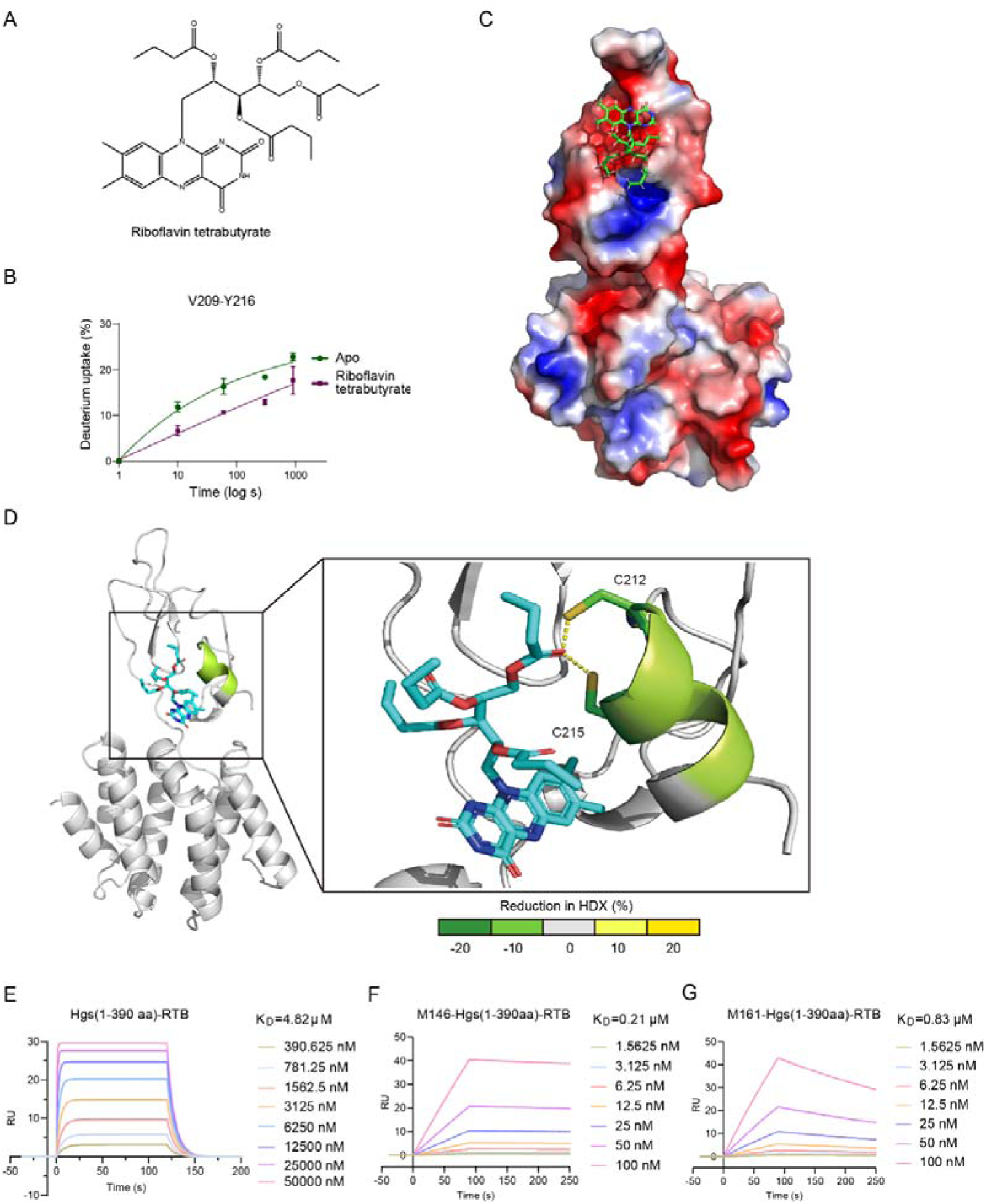
RTB directly targets HGS. (A) The chemical structure of RTB. (B-D) Differential HDX-MS analysis of HGS (1-390) in the presence and absence of RTB is shown as the change in deuterium uptake mapped onto the crystal structure of HGS (PDB: 3ZYQ), highlighting regions affected by RTB binding. The predicted binding pose of RTB with HGS based on the docking result, with RTB represented as indigo sticks (D) and surface representation (C). Deuterium uptake plots for His-tag HGS affected by RTB binding region (green) in the absence (dark green) or presence of RTB (purple), revealing RTB-induced stabilization effects (B). (E-G) SPR analysis of the binding affinity between HGS (1-390) and RTB (E), and binding affinity between HGS (1-390) and peptides M146 (F) or M161 (G) in the presence of RTB. N = 3 independent biological replications. Data are the mean ± SD and analyzed in GraphPad Prism 9.3.

We further explore the direct interaction between HGS and RTB using SPR assay. The results confirmed direct binding between RTB and HGS, with a K_D_ value of 4.82 μM (Fig. 6E). Notably, RTB treatment reduced the binding affinity of HGS for M146 and M161, increasing their K_D_ values from 5.73 nM and 3.84 nM to 0.21 μM and 0.83 μM, respectively (Fig. 6F and 6G).

To investigate whether RTB exerts its anti-coronaviral activity through host factor HGS, the *Hgs*-KO cells were treated with RTB or RDV. HGS ablation significantly reduced coronavirus infection (Fig. S11). Strikingly, RTB treatment in *Hgs*-KO cells failed to inhibit viral infection compared to DMSO-treated *Hgs*-KO cells (Fig. S11). In contrast, RDV maintained potent antiviral activity in *Hgs*-KO cells (Fig. S11). These results demonstrate that RTB’s anti-coronaviral mechanism is HGS-dependent, whereas RDV acts through an HGS-independent pathway.

### RTB inhibits virion assembly

To investigate whether RTB acts at the virion assembly, the effect of RTB on the formation of various coronavirus VLPs was studied. RTB treatment resulted in a concentration-dependent inhibition of VLPs release (Fig. 7A-7D). To confirm these results in cells infected with coronavirus, the effect of RTB was investigated by TEM. TEM observations revealed intracellular mature virions were significantly decreased after treatment of RTB compared with DMSO, while DMVs, the replication organelles, were not significantly changed (Fig. 7E-7H), suggesting that the RTB impedes virion assembly not affects virus replication. Consistent with the effect of HGS deficiency on the M protein localization, the RTB treatment induced M retention in the ER (Fig. 7I and 7J). Collectively, these results demonstrate that RTB prevents M protein trafficking to the ERGIC, thereby inhibiting virion assembly.

**Figure 7.**
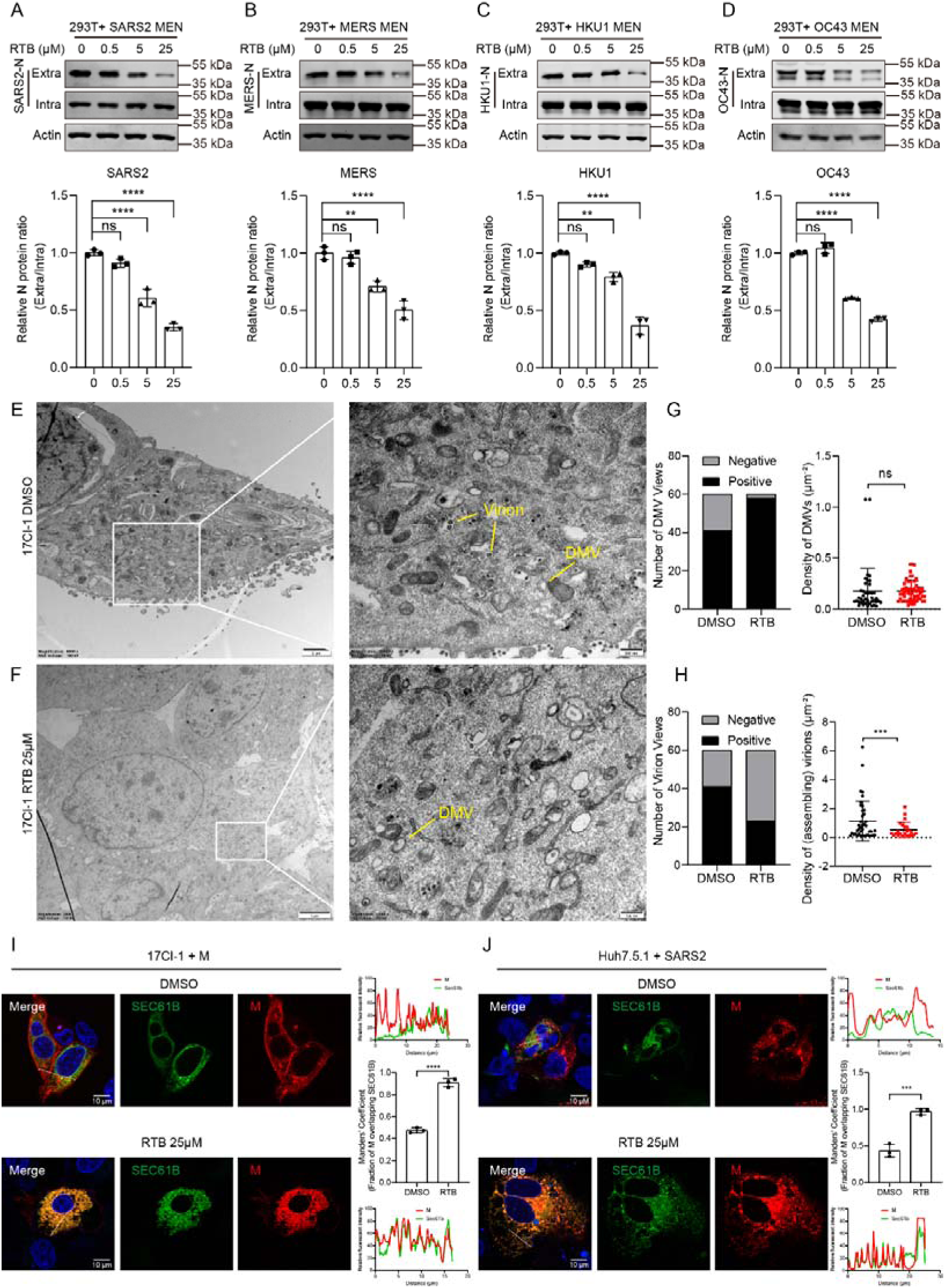
RTB inhibits virion assembly by trapping the M protein in the ER. (A-D) Representative IB analysis of N protein for indicating SARS-CoV-2 (A), MERS (B), HCoV-HKU1 (C) and HCoV-OC43 (D) VLP production in RTB-treated HEK293T cells. HEK293T cells were transfection with VLP system for 48 h. The extracellular secreted protein and intracellular cell lysis were examined by IB with the indicated antibody. Molecular weights are in kDa. N = 3 independent biological replications. (E-H) TEM analysis of DMSO-treated (E) and RTB-treated (25 μM) (F) MHV-infected 17Cl-1 cells (hpi = 24 h, MOI = 1). (G) Quantitative analysis of the number of DMV-positive views (left panel) and the number of DMVs per unit cytoplasmic area (right panel). (H) Quantitative analysis of the number of virion-positive views (left panel) and number of vesicle-contained virions per unit cytoplasmic area (right panel). (I) Representative IF analysis of the co-localization between SEC61B and SARS-CoV-2 M-mCherry protein in DMSO- and RTB-treated (25 μM) 17Cl-1 cells. Scale bar, 10 μm. Quantitative image analysis of M-SEC61B colocalization using Manders’ Coefficient. N = 3 independent biological replications. (J) Representative IF analysis of the co-localization between SEC61B and M protein in SARS-CoV-2 infected Huh7.5.1 cells treatment with or without RTB (25 μM). Scale bar, 10 μm. Quantitative image analysis of M-SEC61B colocalization using Manders’ Coefficient. N = 3 independent biological replications. Data are the mean ± SD. Significance testing for (A-D) was performed with 1-way ANOVA and Tukey’s multiple comparison test. Significance testing for (G-H, I-J) was performed with an unpaired t test. *P ≤ 0.05, **P ≤ 0.005, ***P ≤ 0.0005, ****P ≤ 0.0001, ns, no significance.

Targeted viral assembly inhibitors offer an advantage over polymerase inhibitors in post-infection antiviral therapy and are ideally suited for combination therapies with current protease and polymerase inhibitors. RTB exhibited similar antiviral activity whether treatment began at infection onset or 6 hpi (Fig. S12A). In contrast, the antiviral activity of the polymerase inhibitor RDV decreased in a time-dependent manner when treatment was initiated post-infection (Fig. S12B). Furthermore, RTB demonstrated additive effects *in vitro* when combined with either molnupiravir or nirmatrelvir (Fig. S12C and S12D). RTB combined with molnupiravir also additively decreased viral titers in the lungs of SARS-CoV-2-infected mice (Fig. 5O).

## Discussion

In this study, we utilized a genome-wide CRISPRi screen to identify HGS as a critical host target for pan-coronavirus therapy. We found that HGS directly interacts with M protein, facilitating its trafficking to the ERGIC, a critical step for virion assembly. HGS deficiency or disruption of HGS-M interaction induced M retention in the ER, thereby inhibiting virion assembly. Using a high-throughput screen of >5,000 FDA-approved drugs, we discovered small molecules that bind HGS and disrupt its interaction with the M protein. Among these, RTB exhibits anti-coronavirus activity *in vitro*, in ALI-cultured HBEs and *in vivo*. Collectively, our findings establish HGS as a druggable host target and identify RTB as a promising broad-spectrum antiviral candidate. This work underscores host-directed therapy as a viable strategy against coronaviruses and potentially other viruses.

Targeting virion assembly has emerged as a novel approach for antiviral therapy, with ongoing clinical trials (25). For example, core protein allosteric modulators (CpAMs) have shown efficacy in inducing the assembly of empty and aberrant particles in Hepatitis B virus (26, 27). In the case of HIV-1, peptides derived from the capsid protein have been utilized to hinder polymerization of capsid protein and inhibit virion assembly (28). Subsequently, larger-scale screens for small-molecule were performed to identify the inhibitors of HIV-1 assembly and/or maturation (29). Bevirimat, an inhibitor of HIV-1 assembly and maturation, has shown promising results in clinical trials (30–32). Very recently, two small molecules directly targeting the coronavirus M protein were screened by different groups to inhibit viral infection (17, 18). These molecules bind and stabilize the M protein in its short form, preventing the conformational switch to the long form required for successful virion assembly. However, these compounds exhibited antiviral activity only against a subset of coronaviruses, and resistance emerged in some strains carrying M protein mutations. Consequently, host factors represent promising alternative antiviral targets with potential for broad-spectrum drug development. In this study, we demonstrate that the host factor HGS is essential for pan-coronavirus infection. HGS directly interacts with the M protein, facilitating its trafficking to the ERGIC for virion assembly. Precise M protein localization is critical for assembly. During ER-to-ERGIC trafficking, the M protein conformation converts from the long form to the short form (16). We therefore propose that HGS deficiency traps M in the ER, stabilizing the short form and preventing virion assembly. Targeting the HGS-M interaction thus constitutes a promising antiviral strategy.

Firstly, we designed HGS-targeting peptides derived from the viral M protein to disrupt the HGS-M interaction. These peptides demonstrated broad-spectrum anti-coronavirus activity *in vitro*, in ALI-cultured HBEs, and *in vivo*. This validates viral protein-derived peptides targeting host factors like HGS as a promising antiviral strategy. In the early 1990s, the HIV-1 HR2 domain-derived peptide SJ-2176 could inhibit HIV-1 infection in vitro, leading to the 2003 FDA approved T20 (Fuzeon, enfuvirtide), the first HIV fusion inhibitor-based anti-HIV peptide drug (33). Surprisingly, a HCoV-OC43 HR2-derived peptide, EK1, targeting the HR1 domains of human coronavirus spike proteins, has been confirmed to be effective in inhibiting infection of all circulating human coronaviruses tested, including SARS-CoV, MERS-CoV, HCoV-229E, HCoV-NL63, and HCoV-OC43, as well as 3 bat SARSr-CoVs (34). Our study now establishes a distinct strategy: using viral protein-derived peptides to target essential host factors for antiviral activity.

Furthermore, leveraging structure-activity insights from our peptide inhibitor, we developed a screening strategy to identify FDA-approved small molecules disrupting the HGS-M interaction. This approach identified the vitamin B2 derivative RTB, which demonstrated broad-spectrum anti-coronavirus activity *in vitro*, in ALI-cultured HBEs, and *in vivo*. Crucially, RTB treatment induced M protein retention in the ER, phenocopying the effect of HGS deficiency. This demonstrates that disrupting the HGS-M interaction recapitulates the functional consequences of HGS loss. While targeting an essential host factor like HGS raises potential cytotoxicity concerns, RTB exhibited no cytotoxicity or adverse effects in mice even at high concentrations (500 mg/kg). Moreover, as a derivative of vitamin B2 (riboflavin), RTB has a well-established safety profile from its long-standing use in humans as a dietary supplement. We also provide evidence that RTB directly interacts with HGS and exerts its antiviral effect through this host factor. Although RTB’s antiviral potency is lower than the polymerase inhibitor RDV, it offers distinct advantages: RTB maintains efficacy in post-infection therapy and shows strong potential for combination regimens with current protease and polymerase inhibitors. Future work will focus on determining the structure of the HGS-M complex and developing higher-affinity small molecules targeting HGS.

In addition, our study provides a critical resource for studying coronavirus assembly and egress. Previous studies have predominantly focused on using long-term cytopathic effect-based genome-wide CRISPR-Cas9 screens to identify key host factors involved in coronavirus entry and replication, with a limitation in identifying factors associated with virion assembly and egress (4–8, 35, 36). This is because inhibiting viral entry and replication can enhance cell survival; however, even if virion assembly and egress are obstructed, the virus can still generate cytotoxic viral proteins and induce endo-membrane rearrangements in host cells. Inhibiting virus assembly and egress only hinders the spread of progeny to neighboring cells without alleviating the cytopathic effect within already infected cells. As a result, long-term cytopathic effect-based screens are more likely to identify host factors primarily involved in viral entry and replication rather than in virion assembly and egress. In this study, LAMP1 was introduced as an indicator for coronavirus egress, providing a specific means to identify host factors crucial for virion assembly and egress.

Interestingly, some of these confirmed hits are related to pathways that have also been uncovered in other genome-wide CRISPR screens based on cell survival. Specifically, we found that the rate-limiting enzyme in cholesterol synthesis, HMGCR, is crucial for coronavirus assembly and egress, highlighting the importance of cholesterol homeostasis in virion assembly. Several other genome-wide CRISPR screens based on long-term cytopathic effects have also identified proteins involved in regulating cholesterol homeostasis as key host factors for SARS-CoV-2 and HCoV-OC43 infection (5, 6). Notably, statins, which inhibit HMGCR and thereby reduce cholesterol production, have been associated with improved outcomes in COVID-19 patients (37, 38). Furthermore, genes involved in the biosynthesis and transport of glycosaminoglycans (GAGs), such as EXT1, EXT2, and EXTL3, are essential host factors for carrying out the pan-coronavirus life cycle (6). This aligns with our findings that EXTL2 plays a role in regulating coronavirus assembly and egress. Additionally, some of the identified hits in this study have been shown to regulate infection by other enveloped viruses. For example, INNPL1 (also known as SHIP2) is involved in poxvirus dissemination (39), PDZD8 binds to the HIV-1 Gag polyprotein and enhances HIV-1 infection efficiency (40), and COPA interacts with and regulates trafficking of the SARS-CoV-2 S protein between the ER and Golgi apparatus (41). Overall, this study offers a valuable resource for investigating host factors that play a role in controlling coronavirus assembly and egress.

HGS is a subunit of the ESCRT, which plays a key role in mediating endosomal vesicle budding and the formation of Multivesicular Bodies (MVB) (42). Most enveloped virus families have evolved to utilize ESCRT components, particularly TSG101 and ALIX, to facilitate virus particle assembly, and recruit ESCRT-III and VSP4 complexes for terminal scission events (43). Generally, the structural proteins of enveloped viruses contain specific late domains such as P(S/T)AP, PPXY, or YPXnL (n≤3) that interact with various ESCRT components to facilitate virus assembly and budding (44, 45). However, coronavirus structural proteins do not possess these canonical late domains, raising questions about the involvement of ESCRT components in this process. In this study, it was discovered that HGS interacts directly with and recruits coronavirus M protein to the ERGIC for virion assembly. This is a novel mechanism for virion assembly independent of the late domain signature. Recently, other ESCRT components TSG101 and VPS28 have been demonstrated to play a critical role in membrane budding and terminal scission during coronavirus assembly (46). All these results suggested that ESCRT components participate in coronavirus assembly processes.

## Acknowledgments

The TEM studies were performed at the Center for Biological Imaging (CBI), Institute of Biophysics, Chinese Academy of Sciences. We thank Xixia Li, Zhongshuang Lv, Xueke Tan and Can Peng at CBI for their assistance in EM sample preparations. We thank Junying Jia at the Core Facility, Institute of Biophysics, and Tanghui Liu at the platform of Guangzhou National Laboratory for their assistance in cell sorting. We thank Yan Wang and Jinxin Han at Wuhan University for their assistance in AAV package. This work was supported by the National Natural Science Foundation of China (92469107 to Z.L.; 82170473 to J.S.), Major Project of Guangzhou National Laboratory (GZNL2024A01011 to Y.X.; GZNL2023A01008 to J.S.; SPRG22-002 to Z.L., J.S. and X.C.), National Key R&D Program of China (2024YFA1307400 to Y.X.), Guangdong Province High-level Talent Youth Project (2021QN02Y939 to Z.L.; 2021QN020451 to J.S.).

## Author contributions

X.L. performed the genome-wide CRISPRi screen, most viral experiments and IF. R.C. performed the IB and co-IP, collaborated with K.C. and F.L. to construct the plasmids and KO cell lines. Y.C., Y.L., J.Z. and J.S. carried out FP based high-through screening and HDX analysis. R.B. performed the animal experiments. B.T. carried out the TEM experiments. Y.W. and J.Z. helped with SARS-CoV-2 infection. P.Z. and L.M. helped with WIV1 infection. Y.T. and Q.L. helped with ALI-cultured HBEs. Z.L. initiated the study, directed the research and wrote the manuscript. All the authors discussed the data and reviewed the manuscript.

---

**Figure S1.**
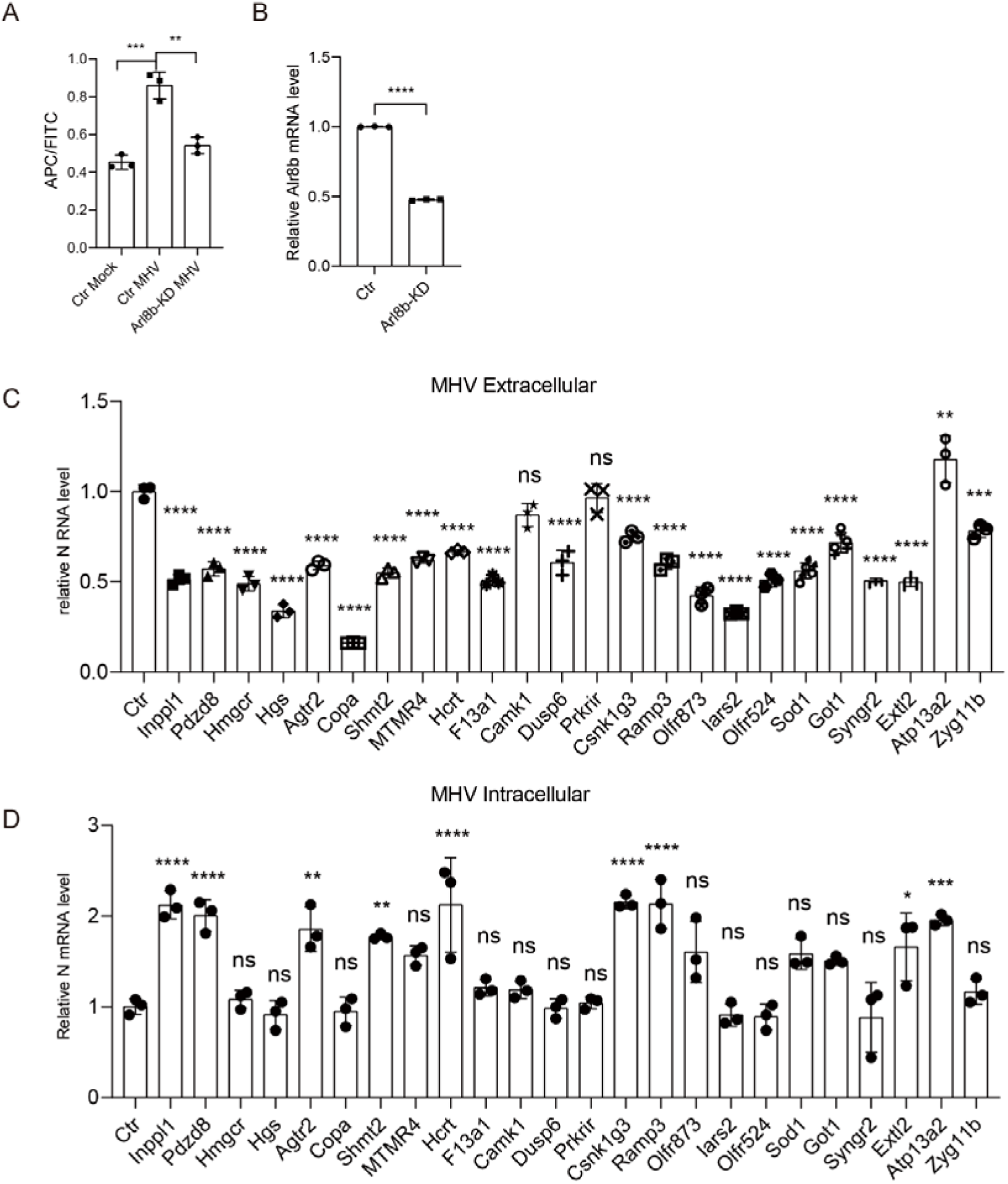
Genome-wide CRISPRi screens identify host factors for pan-coronavirus assembly and egress. (A-B) Flow cytometry analysis of cell surface and total LAMP1 in Ctr and *Arl8b* knockdown 17Cl-1 cells infected with or without MHV at MOI = 1 for 7 h (A). RT-qPCR analysis of *Arl8b* expression in the Ctr and *Arl8b* knockdown 17Cl-1 cells (B). N = 3 independent biological replications. (C-D) Validation of high-confidence host factors. RT-qPCR analysis of extracellular (C) and intracellular (D) viral gRNA levels in dCas9-KRAB expressing 17Cl-1 cells transduced with specific targeted sgRNA after infected with MHV (MOI = 1) for 16 h. N = 3 independent biological replications. Data are the mean ± SD. Significance testing for (A, C and D) was performed with 1-way ANOVA and Tukey’s multiple comparison test. Significance testing for (B) was performed with an unpaired t test. *P ≤ 0.05, **P ≤ 0.005, ***P ≤ 0.0005, ****P ≤ 0.0001, ns, no significance.

**Figure S2.**
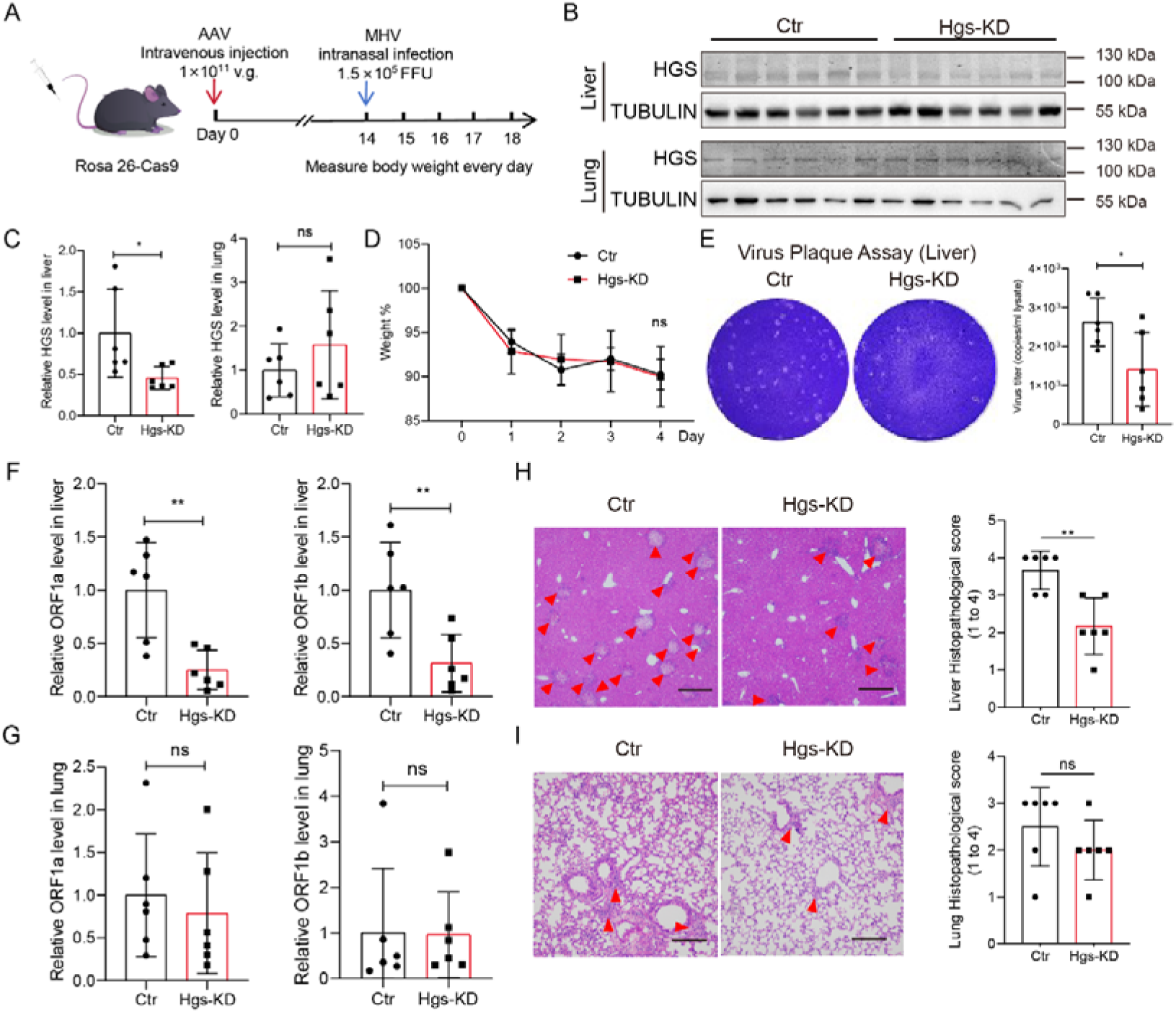
HGS is essential for MHV infection *in vivo*. (A) Schematic illustrating MHV infection in the liver-specific *Hgs* knockdown mice. AAV (1 x 10^11^ v.g.) with scrambled sgRNA or *Hgs* specific targeted sgRNA intravenous injection of Rose26-Cas9 mice at Day 0, and MHV intranasal infection (1.5 x 10^5^ FFU) at Day 14. Measure body weight for 5 days upon infection. N = 6. (B-C) IB analysis of liver and lung HGS expression in the Ctr and *Hgs* knockdown mice (B). Tubulin was used as an internal reference protein. Quantitative analysis of relative HGS protein level in liver and lung (C). N = 6. (D) Body weight loss in the Ctr and *Hgs* knockdown mice post MHV infection for 5 days. N = 6. (E) Viral titration by plaque assay with the supernatant of homogenized liver tissues of the Ctr and *Hgs* knockdown mice on day 4. N = 6. (F-G) RT-qPCR analysis of liver (F) and lung (G) MHV viral ORF1a and ORF1b levels in the Ctr and *Hgs* knockdown mice on day 4. *Gapdh* was used as an internal reference gene. N = 6. (H-I) HE staining analysis of liver (H) and lung (I) tissue in the Ctr and liver-specific *Hgs* knockdown mice. Quantitative analysis of pathological severity scores based on the number of affected area in liver and lung tissues. N = 6. Scale bar = 200 μm Data are the mean ± SD. Significance testing was performed with an unpaired t test. *P ≤ 0.05, **P ≤ 0.005, ***P ≤ 0.0005, ****P ≤ 0.0001, ns, no significance.

**Figure S3.**
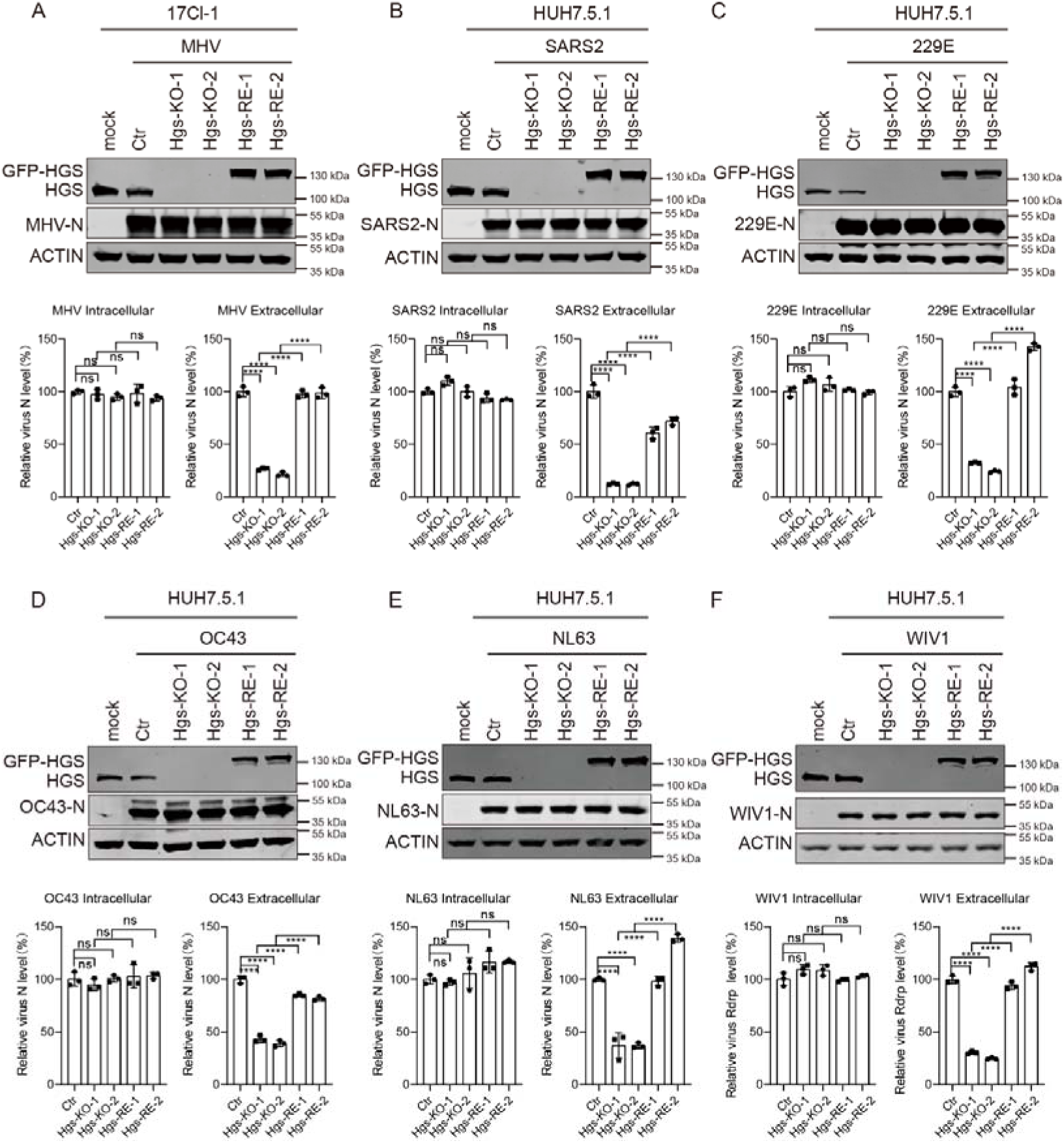
HGS regulates pan-coronavirus assembly and egress. (A-F) RT-qPCR analysis of intracellular (lower left panel) and extracellular (lower right panel) viral gRNA levels in the Ctr, two *Hgs*-KO clones and their respective *Hgs*-recused cells after infected with MHV (17Cl-1 cells for 16 h, Multiplicity of infection, MOI = 1, A), SARS-CoV-2 (Huh7.5.1 cells for 24 h, MOI = 1, B), HCoV-229E (Huh7.5.1 cells for 24 h, MOI = 1, C), HCoV-OC43 (Huh7.5.1 cells for 24 h, MOI = 1, D), HCoV-NL63 (Huh7.5.1 cells for 24 h, MOI = 1, E) and WIV1 (Huh7.5.1 cells for 24 h, MOI = 1, F). IB analysis of HGS and N proteins expression in the Ctr, two *Hgs*-KO clones and their respective *Hgs*-rescued cells (upper panel). Actin was used as an internal reference protein. Molecular weights are in kDa. N = 3 independent biological replications. Data are the mean ± SD. Significance testing for (A-F) was performed with 1-way ANOVA and Tukey’s multiple comparison test. *P ≤ 0.05, **P ≤ 0.005, ***P ≤ 0.0005, ****P ≤ 0.0001, ns, no significance.

**Figure S4.**
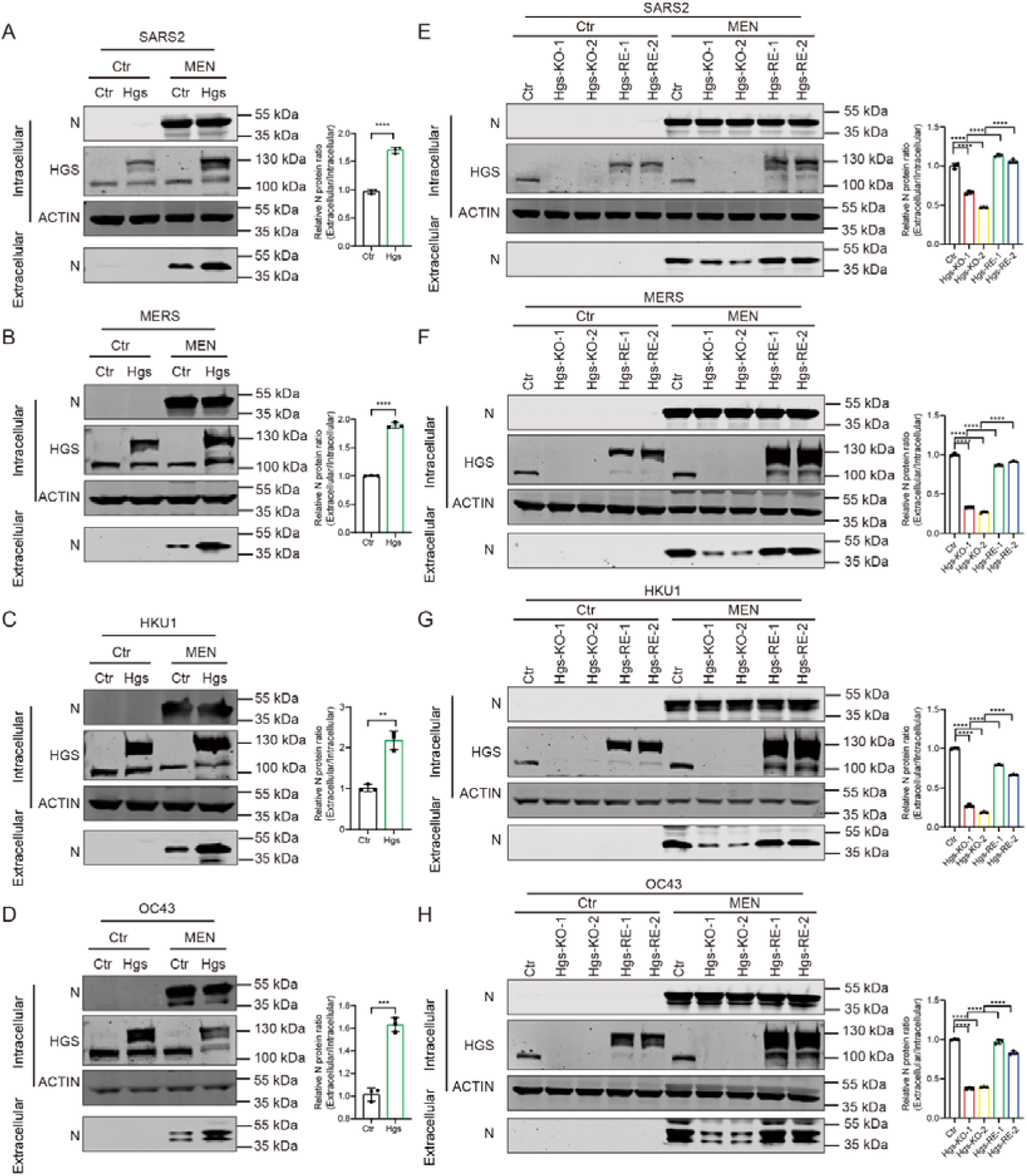
HGS is essential for VLP production. Representative IB analysis of N protein for indicating SARS-CoV-2 (A, E), MERS (B, F), HCoV-HKU1 (C, G) and HCoV-OC43 (D, H) VLP production in the *HGS* overexpression (A-D), two *HGS*-KO clones and their respective *HGS*-recused (E-H) cells. HEK293T cells were transfection with VLP system with HGS-GFP or sgRNA targeted *HGS* for 48 h. The extracellular secreted protein and intracellular cell lysis were examined by IB with the indicated antibody. Molecular weights are in kDa. N = 3 independent biological replications. Data are the mean ± SD. Significance testing for (E-H) was performed with 1-way ANOVA and Tukey’s multiple comparison test. Significance testing for (A-D) was performed with an unpaired t test. *P ≤ 0.05, **P ≤ 0.005, ***P ≤ 0.0005, ****P ≤ 0.0001, ns, no significance.

**Figure S5.**
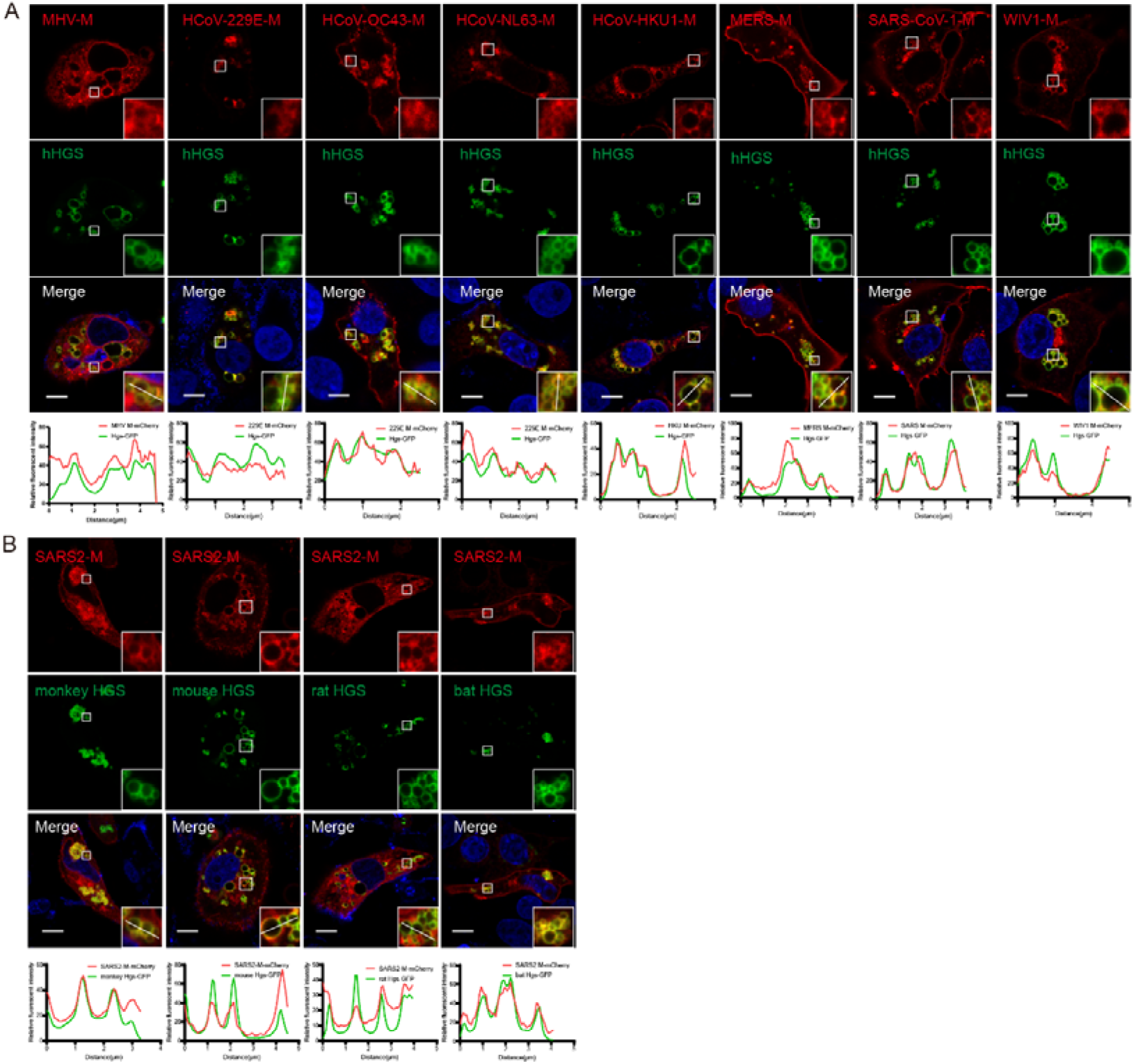
HGS co-localizes with pan-coronavirus M proteins. (A) Representative IF analysis of the co-localization between *Homo sapiens (Human)* HGS and various coronavirus M proteins, including MHV, HCoV-229E, HCoV-OC43, HCoV-NL63, HCoV-HKU1, MERS, SARS-CoV-1, and WIV1. Scale bar, 10 μm. N = 3 independent biological replications. (B) Representative IF analysis of the co-localization between SARS-CoV-2 M protein and different species HGS, including *Chlorocebus sabaeus (Green monkey) (Cercopithecus sabaeus)*, *Rattus norvegicus (Rat)*, *Mus musculus (Mouse)* and *Rhinolophus ferrumequinum (Greater horseshoe bat)*. Scale bar, 10 μm. N = 3 independent biological replications.

**Figure S6.**
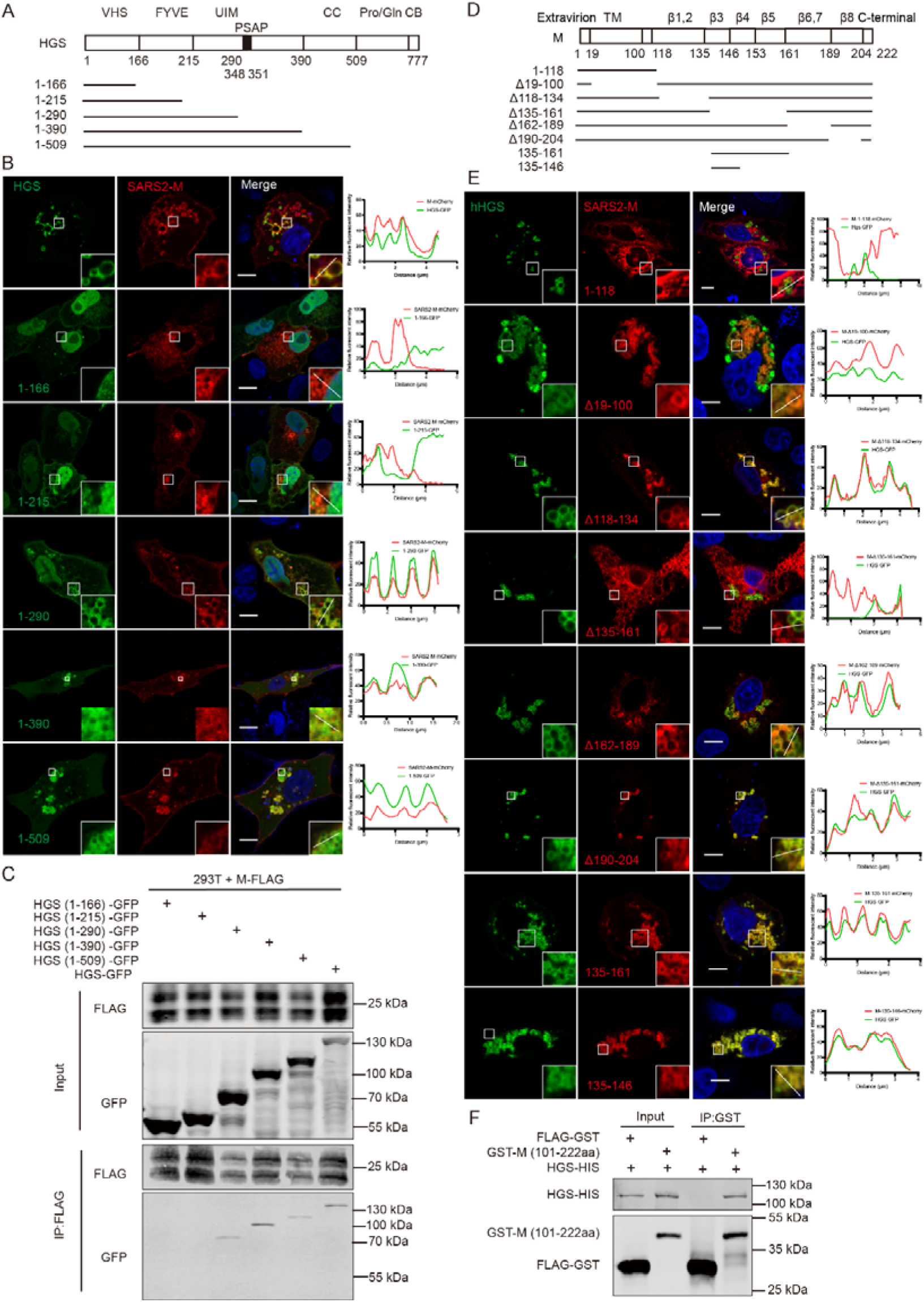
The N-terminal of HGS directly interacts with the intravirion domain of SARS-CoV-2 M protein. (A) Schematic illustrating HGS truncations. (B) Representative IF analysis of the co-localization between M and various truncations of HGS, including HGS (1-166), HGS (1-215), HGS (1-290), HGS (1-390) and HGS (1-509). Scale bar, 10 μm. N = 3 independent experiments. (C) Co-IP analysis of the interaction of M with various truncations of HGS. Proteins were transiently expressed in HEK293T cells, and immunoprecipitates pulled down by FLAG antibody were analyzed by IB with the indicated antibodies. Input represents 5% of the total cell extract used for immunoprecipitation. Molecular weights are in kDa. N = 3 independent biological replications. (D) Schematic illustrating SARS-CoV-2 M truncations. (E) Representative IF analysis of the co-localization between HGS and various truncations of M, including M (1-118), M (Δ19-100), M (Δ118-134), M (Δ135-161), M (Δ162-189), M (Δ190-204), M (135-161) and M (135-146). Scale bar, 10 μm. N = 3 independent biological replications. (F) *In vitro* pull-down analysis of the interaction between HGS and the intravirion domain of M. Purified HGS and the intravirion domain of M proteins were subjected to pull-down assay, and pull-down samples by FLAG antibody were analyzed by IB with the indicated antibodies. FLAG-GST protein was used as a negative control. Input represents 5% of the total proteins used for pull-down. Molecular weights are in kDa. N = 3 independent biological replications.

**Figure S7.**
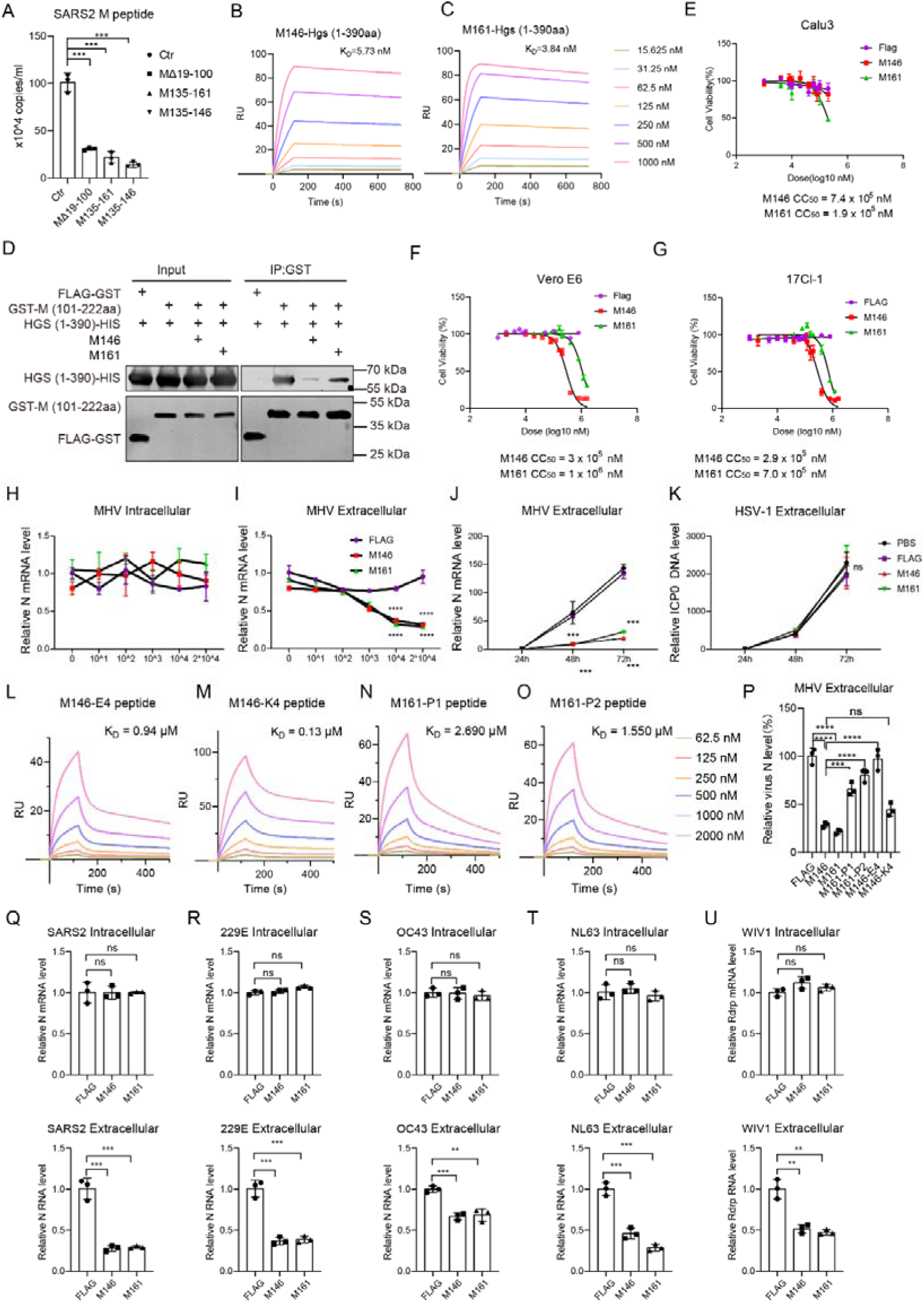
M-derived peptides specifically bind HGS and inhibit pan-coronavirus egress *in vitro*. (A) Plaque assay analysis of extracellular MHV titer levels in the overexpression of SARS-CoV-2 M truncations 17Cl-1 cells. N=3 independent biological replications. (B-C) SPR analysis of the binding affinity between HGS and peptides M146 (B) or M161 (C). N = 3 independent biological replications. (D) *In vitro* pull-down analysis of the competition binding of HGS between peptides M146 and M161 with the intravirion domain of M. Purified HGS (1-390) and the intravirion domain of M proteins with or without peptides M146 or M161 were subjected to pull-down assay, and pull-down samples by GST antibody were analysed by IB with the indicated antibodies. FLAG-GST protein was used as a negative control. Input represents 5% of the total proteins used for pull-down. Molecular weights are in kDa. N = 3 independent biological replications. (E-G) CCK-8 analysis of the cytotoxic effects of M146 and M161 peptides on Vero E6, 17Cl-1 and Calu-3 Cells. N = 3 independent biological replications. (H-I) RT-qPCR analysis of intracellular (H) and extracellular (I) viral gRNA levels in the different doses of FLAG, M146 or M161 peptides treated 17Cl-1 cells after infected with MHV (MOI = 1) for 24 h. N = 3 independent biological replications. (J-K) RT-qPCR analysis of extracellular MHV viral gRNA or HSV-1 viral gDNA levels in the FLAG, M146 or M161 peptides treated (10^4^ nM) 17Cl-1 cells after infected with MHV (MOI = 1) for 24 h, 48 h or 72 h. N = 3 independent biological replications. (L-O) SPR analysis of the binding affinity between HGS and mutated peptides M146-E4 (L), M146-K4 (M), M161-P1 (N) and M161-P2 (O). N = 3 independent biological replications. (P) RT-qPCR analysis of extracellular MHV viral gRNA in the FLAG, M146, M161, M146-E4, M146-K4, M161-P1 and M161-P2 treated (10^4^ nM) 17Cl-1 cells after infected with MHV (MOI = 1) for 24 h. N = 3 independent biological replications. (Q-U) RT-qPCR analysis of intracellular and extracellular SARS-CoV-2 (MOI = 0.5) (Q), HCoV-229E (MOI = 1) (R), HCoV-OC43 (MOI = 1) (S), HCoV-NL63 (MOI = 1) (T) and WIV1 (MOI = 1) (U) viral gRNA levels in infected Vero E6 cells treated with 10^4^ nM FLAG, M146 or M161 peptides for 24 h. N = 3 independent biological replications. Data are the mean ± SD. Significance testing was performed with 1-way ANOVA and Tukey’s multiple comparison test. ***P ≤ 0.0005, ****P ≤ 0.0001, ns, no significance.

**Figure S8.**
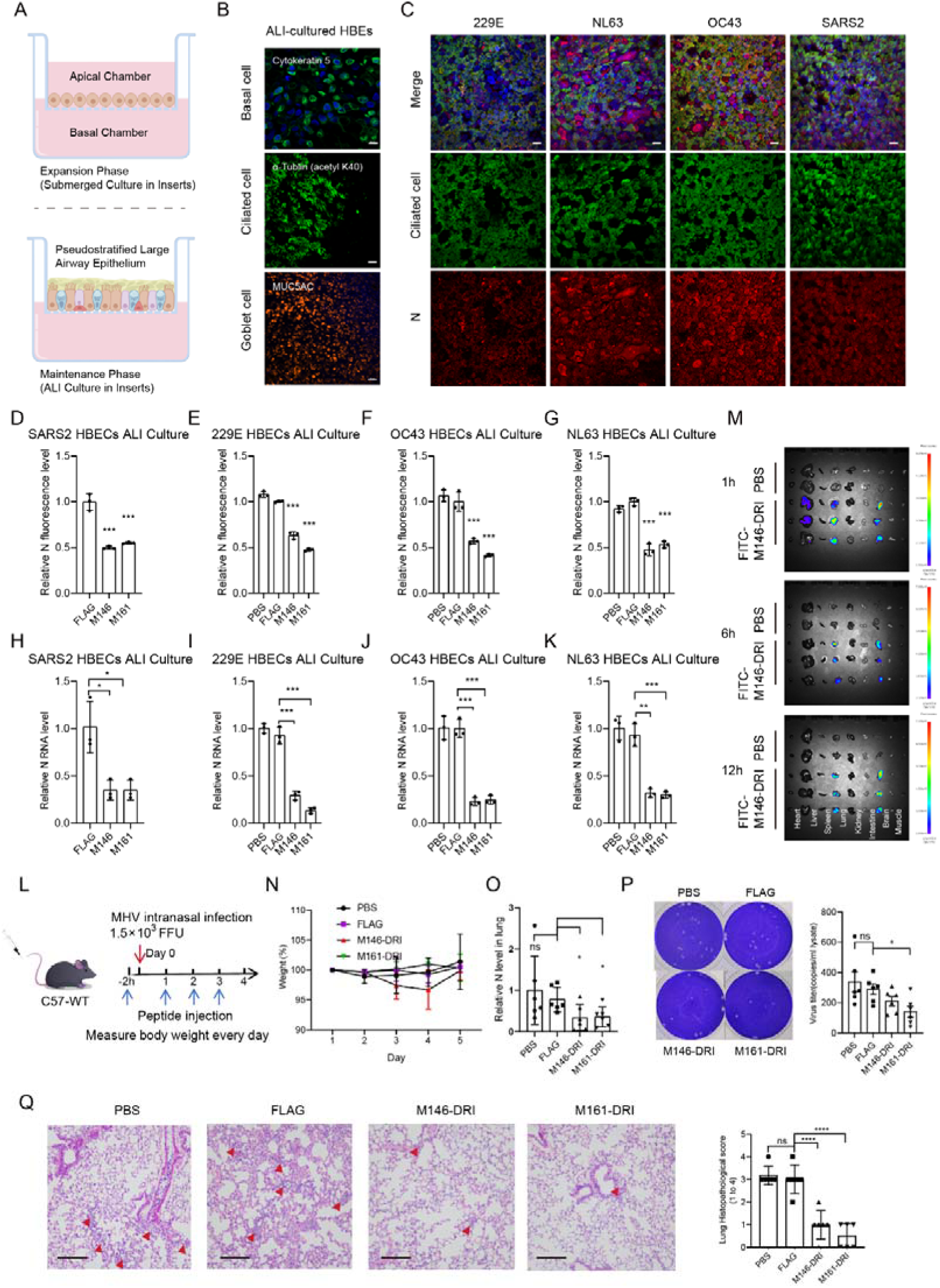
M-derived peptides alleviate the coronavirus infection in ALI-cultured HBEs and *in vivo*. (A) Schematic illustrating primary HBEs in ALI differentiation. (B) Representative IF analysis of basal cell, ciliated cell and goblet cell in the fully differentiated ALI-cultured HBEs. N = 3 independent biological replications. (C) Representative IF analysis of HCoV-229E (MOI = 1, hpi = 96 h), HCoV-NL63 (MOI = 1, hpi = 96 h), HCoV-OC43 (MOI = 1, hpi = 96 h), SARS-CoV-2 (MOI = 0.5, hpi = 96 h) infected ALI-cultured HBEs. N = 3 independent biological replications. (D-G) Quantitative analysis of relative N protein fluorescence levels in the SARS-CoV-2 (MOI = 0.5) (D), HCoV-229E (MOI = 1) (E), HCoV-OC43 (MOI = 1) (F) and HCoV-NL63 (MOI = 1) (G) infected ALI-cultured HBEs treated with 10^4^ nM FLAG, M146 or M161 peptides for 96 h. N = 3 independent biological replications. (H-K) RT-qPCR analysis of extracellular viral gRNA levels in the SARS-CoV-2 (MOI = 0.5) (H), HCoV-229E (MOI = 1) (I), HCoV-OC43 (MOI = 1) (J) and HCoV-NL63 (MOI = 1) (K) infected ALI-cultured HBEs treatment with 10^4^ nM FLAG, M146 or M161 peptides for 96 h. N = 3 independent biological replications. (L) Schematic illustrating MHV-infected mice treatment with HGS-targeted peptides. MHV-infected mice were pre-treated with PBS, FLAG, M161-DRI and M146-DRI peptides (15 mg/kg body weight) for 2 h, afterwards every 24 h for 4 days. (M) *Ex vivo* fluorescence imaging analysis of the distribution of the FITC-labeled M146 peptide in different organs after intravenous injection for 1 h, 6 h, and 12 h, including the heart, liver, spleen, lung, kidney, intestine, brain and muscle. (N) Measure body weight for 5 days. N = 6. (O) RT-qPCR analysis of MHV viral N levels in the lung of the mice treatment with PBS, FLAG, M146-DRI or M161-DRI. *Gapdh* was used as an internal reference gene. N = 6. (P) Viral titration by plaque assay with the supernatant of homogenized lung tissues of the mice treatment with PBS, FLAG, M146-DRI or M161-DRI on day 4. N = 6. (Q) HE staining analysis of lung tissue in the PBS, FLAG, M146-DRI or M161-DRI treated mice. Quantitative analysis of pathological severity scores based on the percentage of affected area in lung tissues. N = 6. Data are the mean ± SD. Significance testing was performed with 1-way ANOVA and Tukey’s multiple comparison test. *P ≤ 0.05, **P ≤ 0.005, ***P ≤ 0.0005, ****P ≤ 0.0001, ns, no significance.

**Figure S9.**
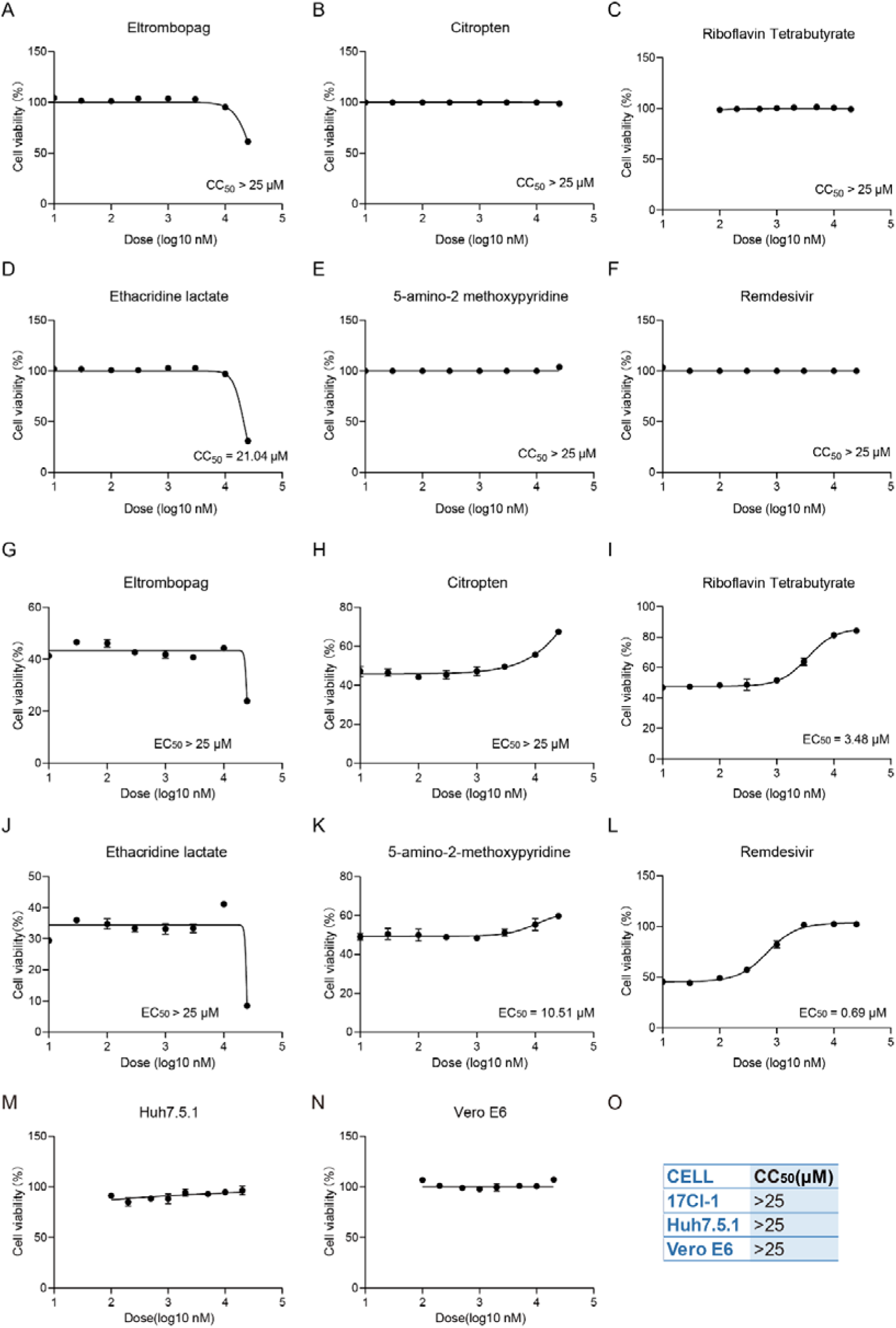
The CC_50_ and EC_50_ of 5 candidate hits. (A-F) CCK-8 analysis of the cytotoxic effects of the top 5 hits on 17Cl-1 (24 h). (G-L) The inhibition of the top 5 hits and the value of EC_50_ is calculated according to MHV-infected 17Cl-1 cell viability by CCK-8 (MOI = 0.1, hpi = 24 h). (M-O) CCK-8 analysis of the cytotoxic effects of RTB on Huh7.5.1 and Vero E6 (24 h). Data were analyzed in GraphPad Prism 9.3.

**Figure S10.**
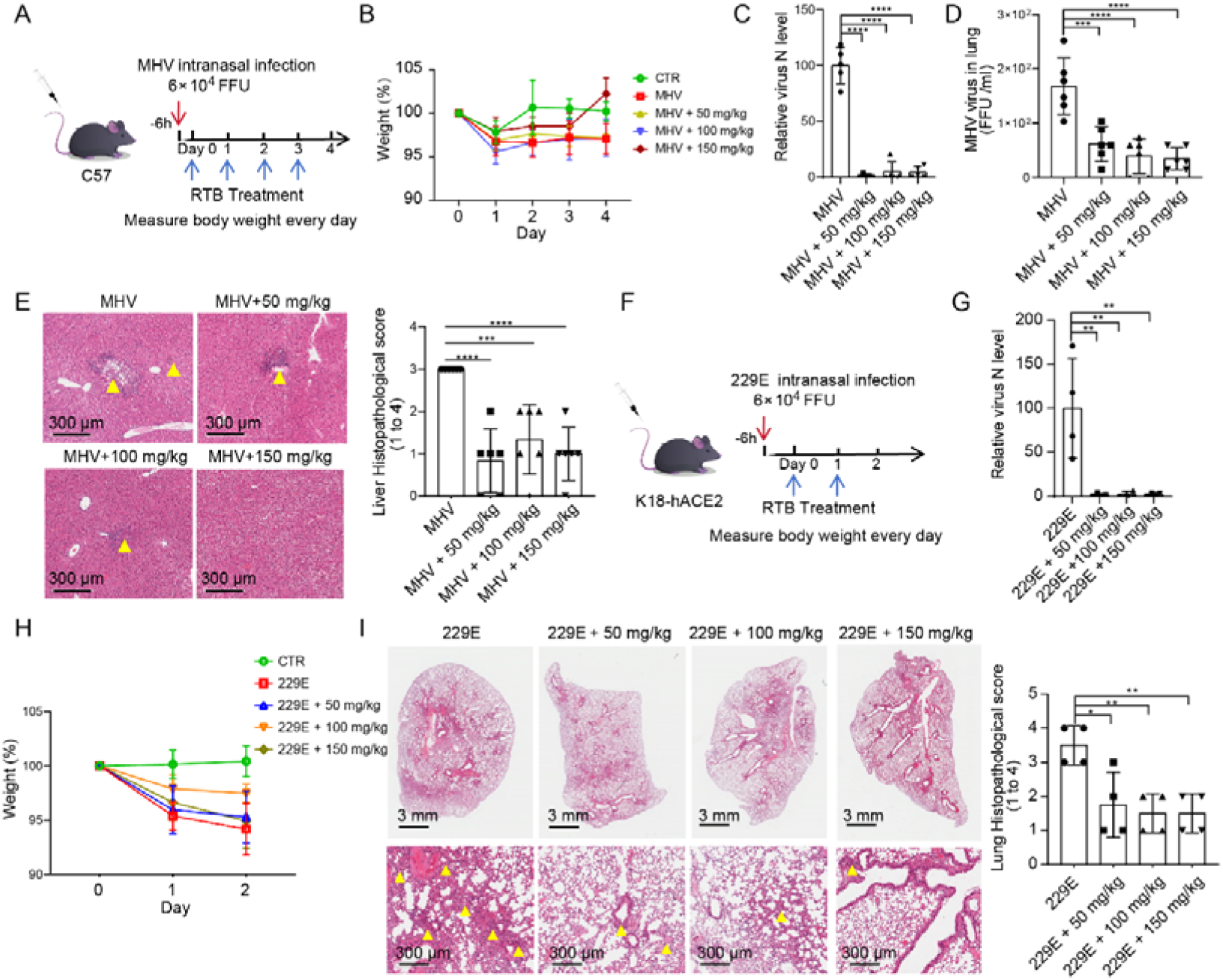
RTB alleviates MHV and HCoV-229E infection *in vivo*. (A-B) Schematic illustrating MHV-infected mice treatment with RTB. MHV-infected mice were treated with PBS and RTB (50, 100, 150 mg/kg body weight) every 24 h for 4 days (A). Measure body weight for 4 days (B). (N = 6 for each group) (C) RT-qPCR analysis of MHV viral N levels in liver of the mice treatment with PBS or RTB. *Gapdh* was used as an internal reference gene. (N = 5 for each group) (D) Viral titration by FFA with the supernatant of homogenized lung tissues on day 4. (N = 6 for each group) (E) HE staining analysis of liver tissue in PBS or RTB treated mice. Quantitative analysis of pathological severity scores based on the number of affected area in liver tissues. N = 9. Scale bar = 300 μm. (F) Schematic illustrating HCoV-229E-infected K18-hACE2 mice treatment with RTB. HCoV-229E-infected mice were treated with PBS and RTB (50, 100, 150 mg/kg body weight) every 24 h for 2 days. (G) RT-qPCR analysis of HCoV-229E viral N levels in lung of the mice treatment with PBS or RTB. *Gapdh* was used as an internal reference gene. (N = 4 for each group) (H) Measure body weight for 2 days. (N = 4 for each group) (I) HE staining analysis of lung tissue in PBS or RTB treated mice. Quantitative analysis of pathological severity scores based on the number of affected area in liver tissues. (N = 4 for each group), Scale bar = 300 μm. Data are the mean ± SD. Significance testing was performed with 1-way ANOVA and Tukey’s multiple comparison test. *P ≤ 0.05, **P ≤ 0.005, ***P ≤ 0.0005, ****P ≤ 0.0001, ns, no significance.

**Figure S11.**
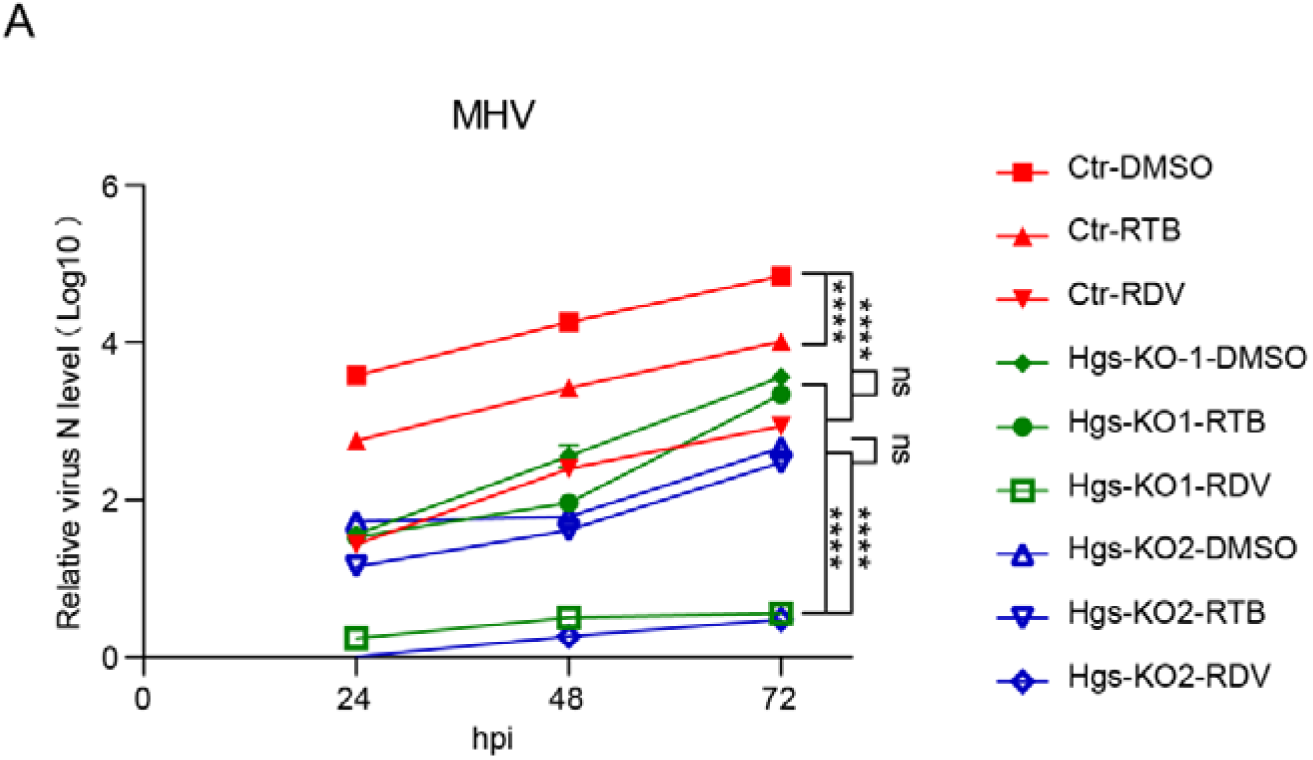
The anti-coronavirus activity of RTB is HGS-dependent. RT-qPCR analysis of extracellular MHV viral gRNA levels in the Ctr, two *Hgs*-KO clones 17Cl-1 cells treated with DMSO, RTB (25 μM) or RDV (5 μM) (hpi = 24 h, 48 h, 96 h respectively, MOI = 1). N = 3 independent biological replications. Data are the mean ± SD. Significance testing was performed with 1-way ANOVA and Tukey’s multiple comparison test. *P ≤ 0.05, **P ≤ 0.005, ***P ≤ 0.0005, ****P ≤ 0.0001, ns, no significance.

**Figure S12.**
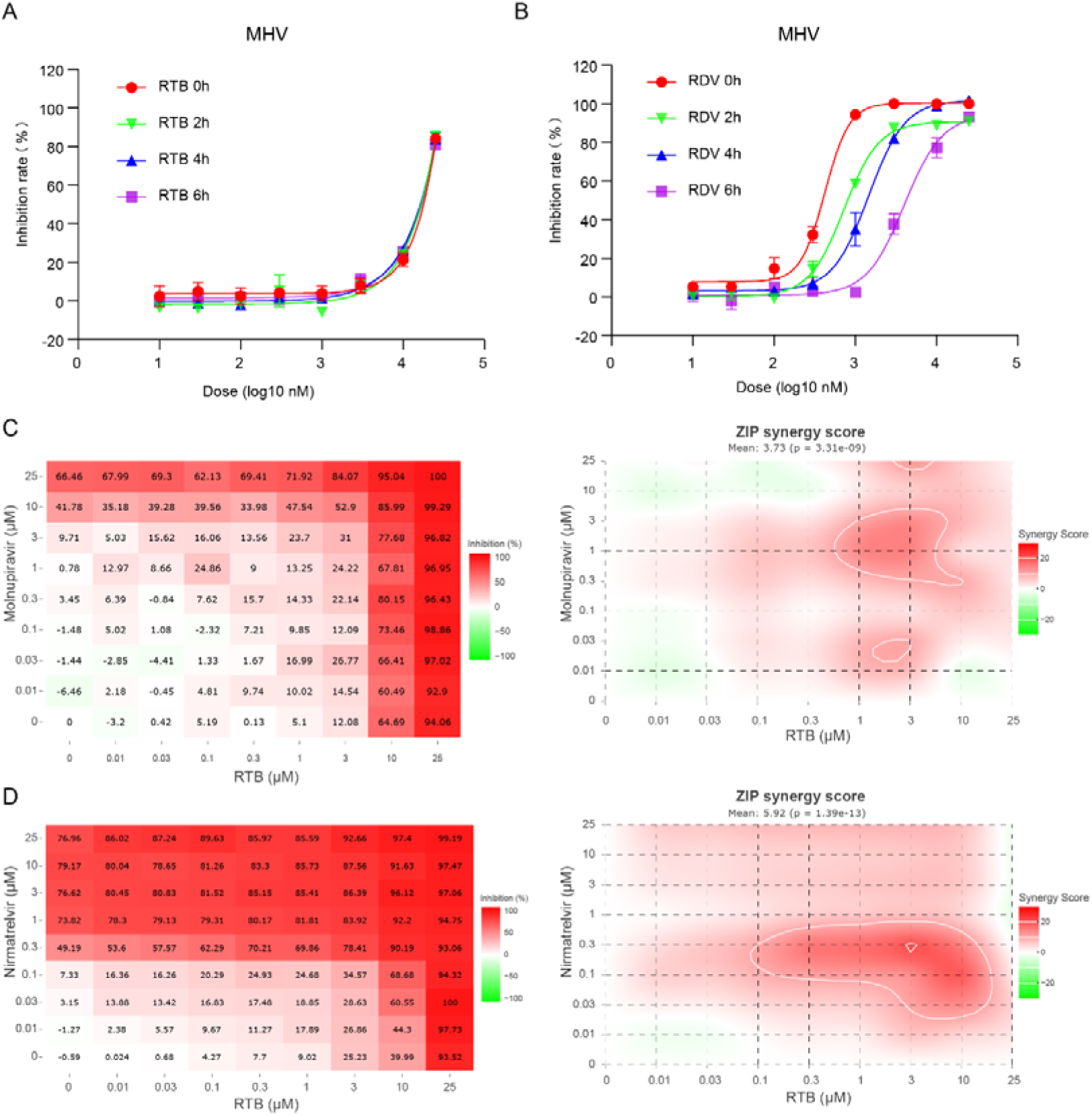
RTB offers an advantage over polymerase inhibitors in post-infection antiviral therapy and is ideally suited for combination therapies with current protease and polymerase inhibitors. (A-B) Time-of-drug-addition assay. The inhibition value of RTB (25 μM) or RDV (5 μM) addition at 0, 2, 4 or 6 hpi is calculated by IFA. (C-D) Combining RTB with Molnupiravir or Nirmatrelvir results in additive antiviral activity in vitro. An antiviral assay was performed by infecting 17Cl-1 cells with MHV and the value virus inhibition rate is calculated by IFA. Left, dose-response matrix for RTB and Molnupiravir (C) or RTB and nirmatrelvir (D) representing average %inhibition of virus replication. Right, heat map of the delta scores (%) for the same combinations and ZIP analysis where δ[=[0, δ[>[0, and δ[<[0 correspond to zero interaction, synergy, and antagonism, respectively. The overall zero interaction potency (ZIP) score represents the response beyond expectation (in %). In the range −10[<[ZIP[<[10, the compounds are likely to act in an additive manner, Score ≥10 indicate synergism. The results shown represent the means of 3 independent experiments for each combination. Data were then analysed with the SynergyFinder webtool based on zero interaction potency (ZIP) model.

