## Supplementary material for "Targeting the host factor HGS-viral membrane protein interaction in coronavirus infection": methods

**Sex as a biological variable.**

Our study examined male and female mice, and we did not observe significant differences between both sexes.

**Cell culture and virus**

HEK293T, 17Cl-1, LLC-MK2, Vero E6, Huh7 and Huh7.5.1 cells were cultured in Dulbecco's Modified Eagle's Medium (DMEM). HRT-18 cells were cultured in RPMI 1640 medium. Calu3 cells were grown in Minimum Essential Medium (MEM). All cells were cultured in indicated medium supplied with 10% fetal bovine serum (FBS) and 1% antibiotics at 37 °C with 5% CO_2_. A list of cell lines used in this study is provided in Table S2.

MHV virus was a gift from Prof. Hongyu Deng (Institute of Biophysics, Chinese Academy of Sciences). HCoV-OC43 (ATCC VR-1558), HCoV-229E (ATCC VR-740), and HCoV-NL63 (ATCC NR-470) were gifted from Prof. Jincun Zhao (Guangzhou National Laboratory). HSV-1 was gifted from Prof. Ersheng Kuang (Zhongshan School of Medicine, Sun Yat-Sen University). WIV1 was provided by Prof. Peng Zhou (Guangzhou National Laboratory). SARS-CoV-2 (Genebank accession no. MT123290.1) was a clinical isolate obtained from the First Affiliated Hospital of Guangzhou Medical University. MHV, HCoV-OC43, HCoV-229E, HCoV-NL63 and SARS-CoV-2 were propagated in 17Cl-1, HRT-18, Huh-7, LLC-MK2, and Vero E6 cells, respectively. Viral stocks were stored at −80 °C. SARS-CoV-2 infection was conducted in Guangzhou Customs District Technology Center BSL-3 Laboratory and Guangzhou National Laboratory BSL-3 Laboratory, while other infections were carried out in BSL-2 laboratory of Guangzhou National Laboratory.

**Genome-wide CRISPRi screen**

The genome-wide CRISPRi screen was modified from a previous report (1). 17Cl-1 stably expressing dCAS9-KRAB cells were transduced with lentiviruses carrying the mCRISPRi-v2 library (MOI = 0.3, provided by Prof. Likun Wang, Institute of Biophysics, Chinese Academy of Sciences) followed by puromycin selection (2 μg/mL) for 5 days to generate library cells. Subsequently, a total of 1 × 10^9^ library cells were infected with MHV at MOI = 1 for 7 h. Following infection, the cells were harvested with accutase (Cat. No. 07920) and stained with APC anti-human CD107a (LAMP-1) antibody (Cat. No. 328620). The cells were washed unbound antibody, fixed with BD Cytofix/Cytoperm™ solution (Cat. No. 554722) and permeabilized by washing in 1 × BD Perm/Wash buffer (Cat. No. 554723). The fixed/permeabilized cells were then suspended in BD Perm/Wash buffer containing FITC anti-human CD107a (LAMP-1) antibody, incubated at 4 °C in the dark for 30 min, and washed before flow cytometric analysis. Cells were filtered through a 40-µm cell strainer before FACS sorting using a BD FACS Aria III. Cell sorting was performed based on the ratio of APC and FITC signals of the cells.

Sorted cells were subjected to extract cellular DNA using the NucleoSpin® Blood (Cat. No. 740951.50) according to the manufacturer’s instruction. SgRNA sequences were amplified using Phanta Max Super-Fidelity DNA Polymerase (Cat. No. P505-d1) and amplicons were purified from 2.5% agarose gel. Amplicons were sequenced using an Illumina Novaseq6000. The sgRNA library (CRISPRi_v2_mouse) was prepared as Comma Separated file containing siRNA id, sequence, and the target gene. The raw sequencing reads (fastq files) were trimmed to remove adaptor by Cutadapt (ver. 4.6). Mapping, counting, enrichment analysis of sgRNA, and gene-level scores were performed using MAGeCK package (ver.0.5.9.5).

**Animals**

Hgs^flox/-^ mice (Strain NO. S-CKO-02885) and K18-hACE2-2A-CreERT2 mice (Strain NO. C001244) in the C57BL/6 background were purchased from Cyagen (Suzhou) Biological Information Technology Co, Ltd. These HGS-floxed mice were then crossed with K18-hACE2-2A-CreERT2 mice to produce double-transgenic offspring: K18-hACE2-2A-CreERT2/HGS-floxed mice. To achieve conditional *Hgs* KO specifically in hACE2-expressing cells, intraperitoneal tamoxifen injections were administered to induce Cre-mediated recombination. All mice were maintained under specific pathogen-free (SPF) conditions and were handled according to the guidelines of the institutional animal guidelines of the animal facilities. Mice were genotyped by PCR amplification of tail DNA using primers specific for Cre recombinase (5′-CATATTGGCAGAACGAAAACGC-3′; 5′-CCTGTTTCACTATCCAGGTTACGG-3′), HGS loxP1 sites (5′-AACCCTTAGAGGGAAACGCAAATC-3′; 5′-TCTAGGGTCCACAGCTAAACTCTC-3′) and HGS loxP2 sites (5′-CTGCTGCAAGGCTACAAGAGT-3′; 5′-ATCCTAATCCATGATCCCCTTTCT-3′).

A 15 mg/mL tamoxifen (Sigma-Aldrich, St. Louis, MO) solution was prepared by dissolving 30 mg tamoxifen in 2 mL corn oil (Sigma-Aldrich). Six-week-old K18-hACE2-2A-CreERT2/Hgs^flox/flox^ mice were intraperitoneally injected with tamoxifen solution (75 mg/kg body weight) once daily for five consecutive days. Following induction for 10 days, mice were intranasally inoculated with 1 × 10^4^ FFU SARS-CoV-2 Omicron BA.5 diluted in 50 μL DMEM. The mice were weighed and visually monitored once daily to score morbidity for 10 days. At 2 days post-infection (Dpi), a group of mice were euthanized and lung organ tissues were sampled for virological and histopathological analyses. Mice were euthanized when they became moribund or died because of difficulty eating or drinking. Randomization and allocation concealment were performed, and the operators of animal experiments were blinded to the group allocation. This work was conducted in the BSL-3 Laboratory of Guangzhou Customs District Technology Center.

The ROSA26-Cas9 mice (Strain NO. S23101012) were purchased from Cyagen (Suzhou) Biological Information Technology Co, Ltd. The C57BL/6JGpt mice were purchased from Zhuhai BesTest Biological Technology Co, Ltd. All animal experiments were approved by the Institutional Animal Care and Use Committee (IACUC) of Guangzhou National Laboratory (GZLab-AUCP-2023-01-A7), and performed in BSL-2 Lab of Guangzhou National Laboratory. The mice well fed in BSL-2 Laboratory for several days to adapt to the environment before performing experiments.

ROSA26-Cas9 mice (6-week-old) were randomly allocated and intravenously injected AAV-U6-mCherry-Control or AAV-U6-mCherry-sgHgs virus particles (1 × 10^12^ vg/ml, 100 μL). After 2 weeks, the mice were inoculated intranasally with MHV (1.5 × 10^5^ FFU). Viral challenge was operated under anesthesia to minimize animal suffering. After infection, the mice were weighed daily. At 5 dpi, mice were euthanized to excise the organs of liver and lung. The collected lung and liver tissues were subjected to examination of viral gRNA levels using RT-qPCR, virus titers by virus plaque assay, and assessment of pathological changes via HE staining.

Eight-week-old C57BL/6JGpt mice were randomly divided into four groups and intranasally inoculated with MHV virus (1.5 × 10^3^ FFU). Two hours prior to MHV infection, each group received an intravenous injection of M-derived peptides at indicated concentration. Subsequently, mice were administered peptides via intravenous injection once daily for three consecutive days. Body weights were monitored daily. At 4 dpi, mice were euthanized, and lungs were collected for examination of viral RNA levels using RT-qPCR, virus titers by virus plaque assay, and assessment of pathological changes via HE staining. For tissue distribution analysis, at 1 h, 6 h, and 12 h post-tail vein injection of 300 μg FITC-peptide, nine organs were excised and imaged using a multimodal animal in vivo imaging system (AniView100, BLT).

Eight-week-old K18-hACE2-2A-CreERT2 C57BL/6JGpt mice were randomly assigned to groups and intranasally inoculated with either SARS-CoV-2 Omicron BA.5 virus (2 × 10⁴ FFU) or HCoV-229E virus (6 × 10⁴ FFU). Concurrently with inoculation, each group began receiving once-daily oral gavage of Riboflavin Tetrabutyrate (RTB) or Molnupiravir at the indicated concentrations. Treatment continued for ten consecutive days for SARS-CoV-2 infection or two consecutive days for HCoV-229E infection. RTB (Selleck, S6441) was dissolved at 10, 20 or 30 mg/mL in 20% anhydrous ethanol and 80% corn oil. Molnupiravir (Selleck, S8969) was dissolved at 10 mg/mL in 5% DMSO and 95% PBS. Mice were monitored daily for weight changes and any clinical signs. A subset of mice was euthanized at 2 dpi for lung collection and subsequent virological and histopathological analyses. Additional mice were euthanized upon becoming moribund or if death occurred due to an inability to eat or drink. Infectious viral lung titers were quantified by fluorescent focus assay (FFA). All SARS-CoV-2 infection work was conducted in the BSL-3 laboratory of Guangzhou National Laboratory, while all HCoV-229E work was performed in the BSL-2 laboratory.

Eight-week-old C57BL/6JGpt mice were randomly divided into five groups and intranasally inoculated with MHV virus (6 × 10^4^ FFU). 6 h post MHV infection, each group received once-daily oral gavage of RTB at the indicated concentrations. Treatment continued for four consecutive days. Mice were monitored daily for weight changes and any clinical signs, and euthanized at 4 dpi for lung collection and subsequent virological and histopathological analyses. All MHV work was performed in the BSL-2 laboratory.

A list of animal models used in this study is provided in Table S2.

**Immunohistochemistry**

The lung tissue induced by tamoxifen and the paraffin-coated sections of the non-induced lung tissues were dewaxed in xylene and hydrated in ethanol. The endogenous peroxidase was blocked using a 3% H_2_O_2_ solution. The antigen was retrieved by boiling in a citric acid solution (pH = 6.0), and non-specific binding sites were blocked with 3% BSA at room temperature for 30 min. The tissues were then incubated with anti-HGS antibody (1: 400, 10390-1-AP, Proteintech) at 4 °C overnight. After washing, the sections were conjugated with a horseradish peroxidase (HRP) antibody (1: 2000, ab205718, Abcam) at room temperature for 50 min. The tissues were developed using 3,3’-diaminobenzidine (DAB) reagent, counterstained with hematoxylin, dehydrated, and mounted.

**Virus infection and determination of viral gRNA levels and virus titers**

For MHV and HSV-1 infection, viruses were added into the 17Cl-1 or Vero E6 culture medium for 2 h at MOI = 1, respectively. Culture medium was replaced by fresh complete medium after 2 times wash of PBS, and infected cells were maintained at 37 °C with 5% CO_2_ for 24 h, 48 h and 72 h to collect infected cell lysate and supernatant for afterwards analysis. For HCoV-OC43, HCoV-229E and HCoV-NL63 infection in Huh7 or Vero E6, after adsorption at 34 °C for 2 h (MOI = 1), the virus inoculum was removed and cells were washed with PBS. Fresh medium was replaced and cells were maintained at 34 °C with 5% CO_2_ for 24 h to collect infected cell lysate and supernatant. For SARS-CoV-2 infection in Huh7 or Vero E6, after adsorption at 37 °C for 2 h (MOI = 0.5), the virus inoculum was removed and cells were washed with PBS. Fresh medium was replaced and cells were maintained at 37 °C with 5% CO_2_ for 24 h to collect infected cell lysate and supernatant.

The virus titration was evaluated by RT-qPCR detection of viral N, RdRp or ICP0 genes. RNA virus MHV, HCoV-OC43, HCoV-229E, HCoV-NL63 and SARS-CoV-2 viral gRNA were extracted from cells or supernatants and reverse-transcribed into cDNA, while DNA virus HSV-1 viral gDNA was extracted from supernatants and analyzed by RT-qPCR. MHV titer was also measured by virus plaque assay.

Briefly, for intracellular mature MHV, HCoV-OC43, HCoV-229E and HCoV-NL63 titer determination, approximately 1 × 10^6^ infected cells were lysed in 500 μL complete medium by liquid nitrogen freeze-thaw pretreatment. 100 μL medium with mature HCoV-OC43, HCoV-229E or HCoV-NL63 virus was collected and then added into cell culture to infect 2 × 10^5^ Huh7.5.1 cells for 6 h. Total RNA was extracted from infected cells using TRIzol (Cat. No. 15596018CN), reverse-transcribed into cDNA using HiScript III 1st Strand cDNA Synthesis Kit (+ gDNA wiper) (Cat. No. R312-02), and analyzed by RT-qPCR. The intracellular mature HCoV-OC43, HCoV-229E and HCoV-NL63 virus titers were measured and represented as the N gene level in the infected cell.

For extracellular mature HCoV-OC43, HCoV-229E and HCoV-NL63 virus, 100 μL medium with mature virus was collected and then added into cell culture to infect 2 × 10^5^ cells, while titers were measured and represented in the same way.

For intracellular virus replication detection, total RNA was extracted from cells using TRIzol, reverse-transcribed into cDNA using HiScript III 1st Strand cDNA Synthesis Kit (+gDNA wiper), and analyzed by RT-qPCR according to the manufacturer's protocol.

For quantification of released extracellular virus, total viral RNA or DNA was isolated from the supernatant and EasyPure® Viral DNA/RNA Kit (Cat. No. ER211) was performed according to the manufacturer's protocol to lyse virus and release DNA/RNA. RNA was extracted, reverse-transcribed into cDNA, while DNA was extracted directly and analyzed by RT-qPCR. All primers used in the article were listed in Table S3.

**Virus Plaque Assay**

For intracellular and extracellular mature MHV titer determination, 100 μL medium with mature virus was collected and then diluted with medium and added into cell culture to infect 2 × 10^5^ 17Cl-1 cells at 37 °C for 2 h. 4 mL of overlay solution was added to each well. After 72 h, the overlay solution was discarded, and crystal violet solution was added. The plates were gently shaken for 2 h, then washed with water to remove the crystal violet stain. The total FFU was determined and represented.

To determine the MHV titer in liver and lung tissue homogenates, the samples were diluted 4-fold with PBS, followed by repeated freezing and thawing in liquid nitrogen. The samples were centrifuged at 10,000 × *g* for 10 min at 4 °C and the supernatant was collected. Next, 100 µL of the supernatant was added to L2 cells in a 12-well plate, with the cells at approximately 90% confluence (L2 cells were kindly provided by Prof. Deyin Guo, Guangzhou National Laboratory), and incubated at 37 °C for 2 h. 4 mL of overlay solution was added to each well. After 72 h, the overlay solution was discarded, and crystal violet solution was added. The plates were gently shaken for 2 h, then washed with water to remove the crystal violet stain. The total FFU of lung and liver tissue was determined and represented.

**Focus forming assay (FFA)**

Vero E6 cells were seeded 2 × 10^4^ cells/well in 96-well plates one day before infection. Lung homogenate was serially diluted and used to inoculate Vero E6 cells at 37 °C for 1 h. Inocula were then removed before adding 125 μL per well 1.6% carboxymethylcellulose warmed to 37 °C. After 24 h, cells were fixed with 4% paraformaldehyde and permeabilized with 0.2% Triton X-100. Cells were then incubated with a rabbit anti-SARS-CoV-2 N protein polyclonal antibody (Cat. No.: 40143-T62, Sino Biological, Inc. Beijing), followed by an HRP-labeled goat anti-rabbit secondary antibody (Cat. No. 109-035-088, Jackson ImmunoResearch Laboratories, Inc. West Grove, PA). The foci were visualized by TrueBlue Peroxidase Substrate (cat. no. 50-78-02, KPL, Gaithersburg, MD), and counted with an ELISPOT reader (Cellular Technology Ltd. Cleveland, OH). Viral titers were calculated as FFU per ml.

**Immunofluorescence Assays (IFA)**

Huh7.5.1, HRT-18, LLC-MK2, 17Cl-1 and Vero E6 cells were seeded 4.5 × 10^4^, 4 × 10^4^, 2.5 × 10^4^, 2 × 10^4^ and 2 × 10^4^ cells/well in 96-well plates one day before infection, separately. Cells were infected with HCoV-229E, HCoV-OC43, HCoV-NL63, MHV， SARS-CoV-2 or WIV1 at MOI = 0.1, respectively. After 24 h or 48 h, cells were fixed with 4% paraformaldehyde and permeabilized with 0.2% Triton X-100. Cells were then incubated with a rabbit anti-229E-N, OC43-N, NL63-N, a human anti-MHV-N or a rabbit anti-SARS-CoV-2 N protein polyclonal antibody, followed by an Alexa Fluor 488-labeled goat anti-rabbit or human secondary antibody. Images of the cells were taken and infection of virus was calculated with the Operetta CLS™ high-content analysis system.

**Plasmids and peptides**

Plasmids containing S protein of SARS-CoV-2 WT and SARS-CoV-1 were generously provided Dr. Shibo Jiang (Fudan University), S protein of SARS-CoV-2 BA4.6 and MERS-CoV were gifted by Dr. Shaobo Wang (Guangzhou National Laboratory). Pan-coronavirus VLP plasmids pCMV-MEN (HCoV-HKU1), pCMV-MEN (SARS-CoV-2), pCMV-MEN (MERS-CoV) and pCMV-MEN (HCoV-OC43) were provided from Prof. Guangxia Gao (Institute of Biophysics, Chinese Academy of Sciences). pcDNA3.1-N (SARS-CoV-2)-flag, pcDNA3.1-M (SARS-CoV-2) and pcDNA3.1-E (SARS-CoV-2)-HA were obtained from Dr. Binbin Ding (Guangzhou National Laboratory). pmCherry-N1-M (HCoV-HKU1), pmCherry-N1-M (SARS-CoV-1), pmCherry-N1-M (MERS-CoV) and pEGFP-N1-Hgs *(Greater horseshoe bat)* were synthesized by Rui Biotech, China. Pan-coronavirus structural protein cDNA fragments (S, M), mutations of S (B1.1.529), mutations of M (SARS-CoV-2) and Hgs of different species or mutations were amplified by Phanta MaxSuper-Fidelity DNA polymerase (Cat. No. P505-d2) and cloned into pLVX-mcherry, pLenti-GFP, pmCherry-N1, pEGFP-N1, pBFP-N1 or pcDNA3.1-flag vectors by ClonExpress II One Step Cloning Kit (Cat. No. C112-02). *Homo sapiens* *HGS* was cloned into the pET28a vector to protein expression and purification. Sequences of all the inserted DNAs were confirmed by sequencing (Rui Biotech, China). All sgRNAs were expressed in the pLG1 or plenti-CRISPR-V2 vector. The peptides (> 95% purity) were synthesized by DGpeptides Co., Ltd., Each peptide, containing 1 mg of powder, was stored at -80 °C. These peptides were dissolved in 1× PBS buffer to a final concentration of 1 mg/mL and diluted as needed. A list of peptides used in this study is provided in Table S4.

**AAV and Lentivirus packaging and infection**

HEK293T cells were co-transfected with pLVX or pLenti lentiviral expression plasmid or pLG1 sgRNA plasmid, and psPAX2 packaging plasmid, and pMD2.G envelope plasmid. The supernatants were harvested at 72 h post‐transfection and concentrated by ultracentrifugation at 100,000 × *g* for 1 h, and then, the pellets were suspended in PBS to prepare 100‐fold viral stocks. After preliminary titration, the lentiviral infections were performed following standard procedures.

The AAV2/9 was used to specifically knockdown *Hgs* in the liver of mice. In brief, sgRNA specific to the mouse *Hgs* or a scrambled control sequence was inserted into pAAV-CMV-mCherry and confirmed via sequencing. The sgRNA sequences used for targeting mouse *Hgs* and the scrambled control were GTACTTGGGTTCGTTCCGGA and ACGGAGGCTAAGCGTCGCAA, respectively. Recombinant AAV was produced by a triple-plasmid transfection system comprising pAAV-PC2 Vector, pHelper Vector and pAAV-CMV-sgRNA-mCherry. The viruses were purified using gradient ultracentrifugation and the virus titers were determined by RT-qPCR.

**Generation of KO or knockdown cell lines**

The KO cell lines were generated via CRISPR-Cas9 or CRISPRi system. sgRNAs targeting the indicated genes were designed by the online tool (<https://chopchop.cbu.uib.no/>). The corresponding DNA was synthesized and cloned into lenti-CRISPR-v2 puro, lenti-CRISPR-v2 BSD vector, or pLG-1 vector. 17Cl-1 and Huh7.5.1 cells were infected with lentivirus collected from HEK293T cells and incubated for 48 h with 8 μg/mL polybrene. Transduced cells were selected with 2 μg/mL puromycin and 30 μg/mL blasticidin (Cat. No. ST551, Cat. No. ST018) for 2 weeks. Single cell colonies were verified by IB analysis and sequencing of the PCR products. The sgRNA sequences were provided in Table S5.

**Co-IP and IB**

For endogenous co-IP, Vero E6 cells (2 × 10^6^ cells) were seeded into a 10 cm dish and infected with SARS-CoV-2 (MOI = 0.5). After 24 h, the total cells were harvested and lysed in 1ml ice-cold IP-lysis buffer (50 mM Tris-HCl pH 7.4, 5 mM EDTA, 40 mM β-sodium glycerophosphate, 30 mM NaF, 1 mM PMSF, 1 mM Na_3_VO_4_, 10% glycerol, 1.0% NP-40, and 150 mM NaCl) in the presence of a protease inhibitor cocktail (Roche) and phosphatase inhibitors. For Co-IP, one 10 cm dish of HEK293T cells was transfected with 10-15 ug plasmid for 48 h, and then, the cells were collected and lysed in 1ml ice-cold NP-40 lysis buffer. Cell lysates were precleaned and incubated with anti-HGS, anti-FLAG or anti-HA antibody and protein A/G agarose beads at 4 °C overnight. After washing five times, the immunoprecipitated complexes were sampled with SDS loading buffer and subjected to IB.

Alternatively, the Pan-coronavirus VLPs in the supernatants were precipitated with a final concentration of 10% (m/v) TCA at 4 °C overnight. After washing two times with acetone and one time with methanol, the protein pellets were dissolved in SDS loading buffer and subjected to IB.

For IB, the samples were separated by SDS-PAGE and transferred to NC membranes. The membranes were blocked in 5% non-fat milk in PBST (1 × PBS with 0.05% tween 20) buffer at room temperature for 1 h and then incubated with primary antibodies at 4 °C overnight, and subsequently incubated with the secondary antibodies (IRDye 680/800, LI-COR Biosciences) at room temperature in the dark for 2 h. The images were visualized and captured by the Odyssey M system (LI-COR Biosciences). Relative protein levels were measured by ImageJ and analyzed by the GraphPad Prism (8.0.1). A list of antibodies used in this study is provided in Table S6.

**Protein puriﬁcation and *in vitro* binding assay**

For recombinant HGS protein, the cDNA of human *HGS* was cloned into pET28a vector with C-terminal 6 × HIS-tag and expressed in *Escherichia coli* BL21 (DE3) with 0.4 mM IPTG at 16 °C for 16 h. Bacterium were collected and lysed by sonication in PBS buffer containing 400 mM NaCl, 20 mM imidazole and protease inhibitors on ice. Supernatants were collected after centrifugation at 12,000 × *g* for 30 min at 4 °C, then was incubated with Ni-NTA agarose beads (Smart-lifesciences) for 2 h at 4 °C. The HIS-tagged HGS was eluted by elution buffer (PBS containing 400 mM NaCl and 250 mM imidazole). The following proteins: GST-FLAG (Cat. No. Ag2329), GST-Spike RBD (Cat. No. Ag30689), GST-M (101-222aa) (Cat. No. Ag30691) were purchased from Proteintech Group, lnc., and M-FLAG protein was gifted from Dr. Xuepeng Wei (Guangzhou National Laboratory).

For in vitro binding assay, anti-FLAG nanobody agarose beads were co-incubated with GST-FLAG or M-FLAG protein and Hgs-6 × HIS protein; Glutathione Sepharose (Cat. No. 17075605) was co-incubated with GST-FLAG or GST-M (101-222 aa) protein and HGS-6 × HIS protein. The beads were overnight at 4 °C, and washed three times with lysis buffer (1 × PBS buffer, contain 400 mM NaCl, 2.5 mM EDTA, 0.2% triton) to remove nonspecifically binding protein. The samples were then subjected to immunoblotting. A list of proteins used in this study is provided in Table S4.

**RT-qPCR**

The extraction of total RNAs was performed using TRIzol reagent (Invitrogen). Subsequently, 1 μg of total RNA per sample was reverse transcribed into cDNA using HiScript II Q Select RT SuperMix for qPCR. RT-qPCR was performed using the TB Green® Premix Ex Taq™ II (Tli RNaseH Plus) (Cat. No. RR820A) and calculated by using the 2^−ΔΔCt^ method with GAPDH as an internal reference gene. The qPCR was conducted with a CFX96 Touch System (CFX96 Touch Real-Time PCR Detection System, Bio-Rad Laboratories (Shanghai) Co., Ltd.). The primer sequences for quantitative measurement were shown in Table S3.

**IF**

Cells infected with virus were fixed with 4% paraformaldehyde at room temperature for 15 min and permeabilized with 0.1% Triton X-100 and 3% BSA in PBS at room temperature for 1 h. Cells were then incubated with specific antibodies at 4 °C overnight, followed by an Alexa Fluor-labeled secondary antibody at room temperature in the dark for 2 h. Images of the cells were taken using a NIKON A1 inverted microscope or Carl Zeiss LSM 980. Manders' Coefficient was calculated using ImageJ with the Coloc 2 plug-in to quantify the fraction of one protein that overlapped with another.

**Primary Human Bronchial Epithelial Cells Air-Liquid interface Culture and virus infection**

Primary human bronchial epithelial cells at passage 3 or earlier were seeded at a density of 40,000 cells per well in Transwell inserts (Costar 6.5 mm Transwell, 0.4 µm Pore Polyester Membrane Inserts，Cat. No. 3470) for expansion culture. The cells were cultured in expansion medium (PneumaCult™-Ex Plus Medium, Cat. No. 05040) until tight junctions were formed. Once tight junctions were established, the expansion medium in the upper chamber was discarded, and the medium in the lower chamber was replaced with air-liquid interface (ALI) differentiation medium (PneumaCult-ALI Medium, Cat. No. 05001). Cells were maintained under ALI conditions, with the medium being changed every two days. The differentiation process was carried out for 21 days, during which cell morphology and differentiation status were monitored. The fully differentiated cells were infected with HCoV-OC43, HCoV-229E, HCoV-NL63 or SARS-CoV-2 (MOI = 1), and fresh medium was changed 1 h post infection. Specific peptide was added into the medium every 24 h. After 96 h post infection, cells in Transwell inserts were washed with 140 μL PBS per insert, and the wash fluid was collected to extract virus RNA followed by RT-qPCR. At the same time, cells in inserts were collected and fixed with 4% paraformaldehyde and permeabilized with 0.1% Triton X-100 and 3% BSA in PBS, then Immunofluorescence Assays was performed to detect N protein and the fluorescence was measured using ImageJ.

**TEM**

WT or *Hgs* KO 17Cl-1 cells were grown on ACLAR films and infected with MHV. At 24 hpi the cells were rinsed with PBS and then fixed with 2.5% glutaraldehyde in PBS overnight. The fixed cells were rinsed with 0.1 M PB 2 times and then double distilled water 2 times, each time for 6 min. They were submerged in an aqueous solution containing 1% osmium tetraoxide and 1.5% potassium ferricyanide for 1.5 h and then rinsed with double distilled water 3 times, each time for 6 min. The samples underwent a dehydration process in which the solution submerging the samples was sequentially replaced by 30%, 50%, 70%, 80%, 90%, 100%, 100% and 100% ethanol solutions, each solution submerging the sample for 6 min. The samples were then infiltrated sequentially with 100% acetone for 6 min 2 times, 3:1 acetone/EMbed 812 resin for 1 h, 1:1 acetone/EMbed 812 resin for 2 h, 1:3 acetone/EMbed 812 resin for 3 h, 100 % EMbed 812 resin for 12 h 2 times. The samples were embedded in EMbed 812 resin and underwent solidification at 45 °C for 12 h and then 60 °C for 48 h. Sectioning was performed with a Leica EM UC6 ultramicrotome. The sections were stained with 2% uranyl acetate for 15 min and 1% lead citrate for 5 min. Electron micrographs were acquired with an FEI Tecnai Spirit 120 kV transmission electron microscope operating at 100 kV accelerating voltage.

**Cytotoxicity assay**

The cytotoxic effects of peptides and compounds on Huh7.5.1, 17Cl-1, Vero E6 and Calu-3 Cells were measured by Cell Counting Kit-8 (CCK-8) assay. Briefly, cell monolayer (1 × 10^4^) grown in 96-well plates was incubated with indicated concentrations of M146, M161 peptides or compounds. After 24 h, the culture medium was replaced with CCK-8 (Cat. No. C0039) solution in fresh medium at 37 °C for 3 h. The absorbance at 450 nm was measured by using a Spectrophotometer Multiskan SkyHigh with Touch Screen (Thermo Fisher, USA) or BioTek Synergy H1 multimode reader (Agilent). The CC_50_ value (Cytotoxic Concentration 50%) of peptides or compounds to cells was calculated by the GraphPad Prism 8.0.1 software.

**In vitro antiviral activity assay**

Huh7.5.1, HRT-18, LLC-MK2, 17Cl-1, and Vero E6 cells were seeded in 96-well plates at densities of 4.5 × 10⁴, 4 × 10⁴, 2.5 × 10⁴, 2 × 10⁴, and 2 × 10⁴ cells/well, respectively, one day prior to infection. Cells were infected with HCoV-229E, HCoV-OC43, HCoV-NL63, MHV, WIV1, or SARS-CoV-2 at an MOI of 0.1. Serially diluted compounds were pre-mixed with viruses, and 100 μL of each mixture was inoculated onto cell monolayers. At 24 or 48 hours post-inoculation, IFA were performed using the Operetta CLS™ high-content analysis system. Compound inhibition and EC₅₀ values were calculated from virus infection rates using GraphPad Prism 8.0.1 software. Data represent one of three independent experiments, each conducted with eight concentration gradients in triplicate wells.

**SPR assay**

SPR assay was performed on a Biacore 8K (Cytiva, USA). M peptides were immobilized in 1 μM on a CM5 sensor chip to reached at least 700 resonance units (RU), followed by flowing of hHGS (1-390)-6 × HIS protein at varying concentrations in 1 × PBS (pH 7.4). Binding studies were performed by passing two-fold serial dilutions of purified hHGS (1-390)-6 × HIS protein and measured by a multiple-cycle method with 120 s association and 600 s dissociation at a flow rate of 30 μL/min. For competitive inhibition experiments, the concentration of RTB is 200 μM. In addition, the direct interaction between hHGS (1-390)-6 × HIS protein and RTB was detected using CM7. The resulting SPR sensorgrams were recorded and analyzed using the software provided by the vendors to extract the association and dissociation rate constants (ka and kd) and the binding affinity (K_D_). Each SPR experiment was repeated at least three times and analyzed by the software Prism 8.0.1 software.

**Fluorescence polarization HGS-peptide interaction assays**

Fluorescence polarization assays were performed in black 384-well microplates (Corning, no. 3575) with 20 μL of the assay solution per well, which contains 0.8 μM hHgs (1-390) protein and 180 nM FITC-peptide tracer. The optimal FP buffer was 50 mM HEPES, pH 7.2, 10 mM DTT, 0.1 mM EDTA, pH 7.2 with additives (0.005% Tween-20). Ligand stocks were prepared through a serial dilution in DMSO and added to the wells. Assay mixture was incubated for 1h at room temperature before read plate. All data were collected on a microplate reader (BioTek, SYNERGY H1). Excitation (485/20 nm) and emission (520/20 nm) filters were used for the FP measurements. All the experiments were independently repeated at least three times. The IC_50_ curves of compounds were analyzed in GraphPad Prism 8.3.

**High-throughput screening of an in-house compound library**

In the FP-based high-throughput screening assay, the compounds (10 mM in DMSO stock for a library of 5000+ compounds) were transferred into each well of 384-well that contains 0.8 μM HGS (1-390) protein and 180 nM FITC-peptide tracer using pintool at 33 μM final concentration. In each assay plate, the DMSO and unlabeled peptides were used as negative and positive controls, respectively. Free tracer wells (180 nM tracer only) were also set up as gain value in each assay plate. After incubation for 1 h at RT, the FP was measured with an excitation at 485/20 nm and emission at 520/20 nm for FITC. The top 50 hit compounds were validated with FITC-peptide tracer in a FP assay, each tested in three independent replicates.

**Hydrogen-deuterium exchange (HDX)**

*Hydrogen-deuterium exchange (HDX) mass spectrometry (MS) for peptide identification.* Peptides were identified using tandem MS (MS/MS) with a Q Exactive HF mass spectrometer (Thermo Fisher). Product ion spectra were acquired in data-dependent mode with the top eight most abundant ions selected for the product ion analysis per scan event. The MS/MS data files were entered into pFind for high-confidence peptide identification.

*HDX-MS analysis.* Ten microliters of hHGS (50 mM HEPES, pH 7.5; 50 mM NaCl) was incubated with and without the RTB at a 1: 50 molar ratio (protein: ligand) for 0.5 h before the HDX reacted at 4 °C. Four microliters of protein/protein complex with ligand/peptide was diluted into 16 µL of D_2_O in exchange buffer (50 mM HEPES, pH 7.5; 50 mM NaCl) and incubated for various HDX time points (e.g., 0, 60, 300s) at 4 °C and quenched by mixing with 20 µL of ice-cold 3M guanidine hydrochloride and 1% trifluoroacetic acid. Each quenched sample was immediately injected into the LEAP Pal 3.0 HDX platform. Upon injection, samples were passed through an immobilized pepsin column (2 mm × 2 cm) at 120 µL/min, and the digested peptides were captured on a C18 PepMap300 trap column (Thermo Fisher) and desalted. Peptides were separated with a 2.1 mm x 5 cm C18 separating column (1.9 μm Hypersil Gold, Thermo Fisher) with a linear gradient of 4-40% CH3CN and 0.3% formic acid over 6 min. Sample handling, protein digestion, and peptide separation were conducted at 4 °C. Mass spectrometric data were acquired using a Q Exactive HF mass spectrometer (Thermo Fisher) with a measured resolving power of 65,000 at m/z 400. HDX analyses were performed in triplicate for each preparation of a single protein-ligand complex. The intensity weighted mean m/z centroid value of each peptide envelope was calculated and subsequently converted into a percentage of deuterium incorporation. Statistical significance for the differential HDX data is determined by an unpaired t test for each time point, a procedure that is integrated into the HDX Workbench software (2). Corrections for back exchange were made on the basis of an estimated 70% deuterium recovery and accounting for the known 80% deuterium content of the deuterium exchange buffer.

*Data rendering.* The HDX data from all overlapping peptides were consolidated to individual amino acid values using a residue averaging approach. Briefly, for each residue, the deuterium incorporation values and peptide lengths from all overlapping peptides were assembled. A weighting function was applied in which shorter peptides were weighted more heavily and longer peptides were weighted less heavily. Each of the weighted deuterium incorporation values were then averaged to produce a single value for each amino acid. The initial two residues of each peptide, as well as proline residues, were omitted from the calculations.

**Statistics.**

The data were obtained from at least three independent experiments (n ≥ 3), and the representative data are presented as the mean ± SD as indicated and were corrected for multiple comparisons. ANOVA and Student’s t tests were used to analyze differences in mean values. GraphPad Prism 8.0.1 software was used to calculate the P‐values, and significance is depicted with asterisks as follows: *P ≤ 0.05, **P ≤ 0.005, ***P ≤ 0.0005, ****P ≤ 0.0001.

**Study approval.**

All animal studies were approved by the Guangzhou National Laboratory Animal Care and Use Committees and met stipulations of the NIH *Guide for the Care and Use of Laboratory Animals* (National Academies Press, 2011).

**Data availability.**

The data supporting the findings of this study are documented within the paper and are available from the corresponding authors upon request. Correspondence and requests for materials should be addressed to: Zonghong Li.

1.	Horlbeck MA, Gilbert LA, Villalta JE, Adamson B, Pak RA, Chen Y, et al. Compact and highly active next-generation libraries for CRISPR-mediated gene repression and activation. *eLife.* 2016;5.

2.	Pascal BD, Willis S, Lauer JL, Landgraf RR, West GM, Marciano D, et al. HDX workbench: software for the analysis of H/D exchange MS data. *Journal of the American Society for Mass Spectrometry.* 2012;23(9):1512-21.

**Table S2.** List of Cell lines and animals.

| Human: HEK293T | ATCC | Cat# CRL-3216 |
| --- | --- | --- |
| Human: Huh7 | Cas9X^TM^ | Cat# TCH-C217 |
| Human: Huh7.5.1 | Gifted by Xiancai Ma | N/A |
| Mouse: 17Cl-1 | Gifted by Hongyu Deng | N/A |
| Human: HRT-18 | Gifted by Jincun Zhao | N/A |
| Monkey: LLC-MK2 | Gifted by Jincun Zhao | N/A |
| African green monkey: Vero E6 | Gifted by Xiancai Ma | N/A |
| Human: Calu-3 | Gifted by Xiancai Ma | N/A |
| Mouse: 17Cl-1 *Hgs^KO-1^* | This paper | N/A |
| Mouse: 17Cl-1 *Hgs^KO-2^* | This paper | N/A |
| Mouse: 17Cl-1 *Hgs^RE-1^* | This paper | N/A |
| Mouse: 17Cl-1 *Hgs^RE-2^* | This paper | N/A |
| Human: Huh7.5.1 *Hgs^KO-1^* | This paper | N/A |
| Human: Huh7.5.1 *Hgs^KO-2^* | This paper | N/A |
| Human: Huh7.5.1 *Hgs^RE-1^* | This paper | N/A |
| Human: Huh7.5.1 *Hgs^RE-2^* | This paper | N/A |
| Human: HEK293T *Hgs^KO-1^* | This paper | N/A |
| Human: HEK293T *Hgs^KO-2^* | This paper | N/A |
| Human: HEK293T *Hgs^RE-1^* | This paper | N/A |
| Human: HEK293T *Hgs^RE-2^* | This paper | N/A |
| Mouse: ROSA26-Cas9 | Saiye (Suzhou) Biological Information Technology Co,.Ltd | Cat# S23101012 |
| Mouse: C57BL/6 | Zhuhai BesTest Biological Technology Co,.Ltd | Cat# BST-111 |
| Mouse: K18-hACE2-2A-CreERT2 | Saiye (Suzhou) Biological Information Technology Co,.Ltd | Cat# C001244 |
| Mouse: Hgs (flox/+) C57BL/6 | Saiye (Suzhou) Biological Information Technology Co,.Ltd | Cat# S-CKO-02885 |
| HBECs | Lonza | Cat# CC-2540S |

**Table S3.** List of PCR primers.

| **Gene** | **Sequence (5′ >>> 3′)** | |
| --- | --- | --- |
| Mouse *Arl8b* | Forward | GATACCCACAGTGGGCTTCAAC |
|  | Reverse | TGCATTGACTCCTCGGCAGTAC |
| WIV1 *RdRp* | Forward | GGTCATGTGTGGCGGCTC |
|  | Reverse | GCTGTAACAGCTTGACAAATGTTAAAG |
| nCoV *N* | Forward | CACATTGGCACCCGCAATC |
|  | Reverse | GAGGAACGAGAAGAGGCTTG |
| OC43 *N* | Forward | GGGACCCAAGTAGCGATGAG |
|  | Reverse | TGGCTCTACTACGCGATCCT |
| 229E *N* | Forward | CGCAAGAATTCAGAACCAGAG |
|  | Reverse | GGGAGTCAGGTTCTTCAACAA |
| NL63 *N* | Forward | AGGACCTTAAATTCAGACAACGTTCT |
|  | Reverse | GATTACGTTTGCGATTACCAAGACT |
| MHV *N* | Forward | TGGAAGGTCTGCACCTGCTA |
|  | Reverse | TTTGGCCCACGGGATTG |
| MHV *ORF1a* | Forward | GTTCAGTCTGCCATAATCCG |
|  | Reverse | GTCACAAGAGCACTCCCTG |
| MHV *ORF1b* | Forward | GAGTATGCCTCCAACTCTGC |
|  | Reverse | ACAAACGGAAGCACCACC |
| HSV-1 *ICP0* | Forward | CCCACTATCAGGTACACCAGCTT |
|  | Reverse | CTGCGCTGCGACACCTT |
| Human *GAPDH* | Forward | GTCTCCTCTGACTTCAACAGCG |
|  | Reverse | ACCACCCTGTTGCTGTAGCCAA |
| Mouse *GAPDH* | Forward | CATCACTGCCACCCAGAAGACTG |
|  | Reverse | ATGCCAGTGAGCTTCCCGTTCAG |
| *Cre* recombinase | Forward | CATATTGGCAGAACGAAAACGC |
|  | Reverse | CCTGTTTCACTATCCAGGTTACGG |
| *HGS* loxP1 | Forward | AACCCTTAGAGGGAAACGCAAATC |
|  | Reverse | TCTAGGGTCCACAGCTAAACTCTC |
| *HGS* loxP2 | Forward | CTGCTGCAAGGCTACAAGAGT |
|  | Reverse | ATCCTAATCCATGATCCCCTTTCT |

**Table S4.** List of Chemicals, compounds, peptides, and recombinant proteins.

| Hoechst 33342 | Thermo Fisher | Cat# H1399 |
| --- | --- | --- |
| Puromycin Dihydrochloride | Beyotime Biotechnology | Cat# ST551 |
| Blasticidin S HCl | Beyotime Biotechnology | Cat# ST018 |
| Hygromycin B | Beyotime Biotechnology | Cat# ST1389 |
| Polybrene (Hexadimethrine Bromide) | Beyotime Biotechnology | Cat# C0351 |
| Tamoxifen | Sigma-Aldrich | Cat# 10540-29-1 |
| Protein A/G Agarose Beads | Proteintech | Cat# PR40025 |
| Anti FLAG Nanobody Magarose Beads | KTSM-Life | Cat# KTSM1338 |
| Glutathione Sepharose (GST Tag) | Cytiva | Cat# 17075605 |
| Lipofectamine™ 3000 Transfection Reagent | Thermo Fisher | Cat# L3000015 |
| TRIzol™ Reagent | Invitrogen | Cat# 15596018 |
| Fetal Bovine Serum, Premium Plus | Thermo Fisher | Cat# A5669701 |
| TrypLE™ Express Enzyme (1 × ), phenol red | Thermo Fisher | Cat# 12605028 |
| ACCUTASE™ | STEMCELL Technologies | Cat# 07920 |
| Opti-MEM® I Reduced Serum Medium | Gibco | Cat# 31985070 |
| Penicillin-Streptomycin, Liquid (100 × ) | Invitrogen | Cat# 15140122 |
| EMS Aqueous Glutaraldehyde EM Grade 25% | Electron Microscopy Sciences | Cat# 16220 |
| Peptide FLAG:  GRKKRRQRRRPPQDYKDDDDK | DGpeptides Co., Ltd | N/A |
| Membrane (M) peptide M146 :  GRKKRRQRRRPPQESELVIGAVILR | DGpeptides Co., Ltd | N/A |
| Membrane (M) peptide M146-DRI:  RLIVAGIVLESEQPPRRRQRRKKRG | DGpeptides Co., Ltd | N/A |
| FITC-Membrane (M) peptide M146-DRI:  FITC-RLIVAGIVLESEQPPRRRQRRKKRG | DGpeptides Co., Ltd | N/A |
| Membrane (M) peptide M161:  GRKKRRQRRRPPQESELVIGAVILRGHLRIAG | DGpeptides Co., Ltd | N/A |
| Membrane (M) peptide M161-DRI:  GAIRLHGRLIVAGIVLESEQPPRRRQRRKKRG | DGpeptides Co., Ltd | N/A |
| Membrane (M) M146-E4：  GRKKRRQRRRPPQESELVIGAVILREEEE | DGpeptides Co., Ltd | N/A |
| Membrane (M) M146-K4：  GRKKRRQRRRPPQESELVIGAVILRKKKK | DGpeptides Co., Ltd | N/A |
| Membrane (M) M161-P1：  GRKKRRQRRRPPQESELPIGAVILRGHLRIAG | DGpeptides Co., Ltd | N/A |
| Membrane (M) M161-P2：  GRKKRRQRRRPPQESELPPGAVILRGHLRIAG | DGpeptides Co., Ltd | N/A |
| DYKDDDDK tag Fusion protein (FLAG-GST) | Proteintech | Cat# Ag2329 |
| 2019-nCOV Spike protein (RBD) Fusion Protein (GST-RBD) | Proteintech | Cat# Ag30689 |
| Membrane glycoprotein Fusion Protein (GST-M :101-222 aa) | Proteintech | Cat# Ag30691 |
| Immobilon NC Transfer Membrane | Merck millipore | Cat# HATF00010 |
| Peptide FLAG-M | This paper | N/A |
| Peptide HIS-HGS | This paper | N/A |
| Peptide HIS-HGS(1-390) | This paper | N/A |
| Riboflavin Tetrabutyrate | Selleck | S6441 |
| Nirmatrelvir (PF-07321332) | Selleck | S9866 |
| Molnupiravir (EIDD-2801) | Selleck | S8969 |
| Remdesivir (GS-5734) | TargetMol | T7766 |

**Table S5.** List of sgRNA primer sequence.

| **Name** | **Sequence (5′ to 3′)** |
| --- | --- |
| Mouse *Inppl1* target (KD) | ACCCGAGTCGCCCTAGAGG |
| Mouse *Pdzd8* target (KD) | CCGCTCGGGCCGCCCCCGG |
| Mouse *Hmgcr* target (KD) | GCGTCCGCCAGCTCACCTC |
| Mouse *Hgs* target (KD) | CTGCAGCGTCGGTCCGGAG |
| Mouse *Agtr2* target (KD) | CTCAGAGGCTGGCGATGGA |
| Mouse *Copa* target (KD) | GAAGCCTTGGGAGTGCCAC |
| Mouse *Shmt2* target (KD) | CTAGCCACTAACTCTGTAT |
| Mouse *Mtmr4* target (KD) | CAGCTGCGGGAGACGGAAG |
| Mouse *Hcrt* target (KD) | AGCACTGAGAGAGGAGTAT |
| Mouse *F13a1* target (KD) | GGCAAAGTGACCAGAAAGT |
| Mouse *Camk1* target (KD) | CTGGCTGCGAGGGCCGAGG |
| Mouse *Dusp6* target (KD) | ACTGGGTAGGAACAAAACT |
| Mouse *Prkrir* target (KD) | TCCCCTCCCCGGCCCCGCG |
| Mouse *Csnk1g3* target (KD) | CCGGAAGTGAACACATGAG |
| Mouse *Ramp3* target (KD) | CAGCAACAGCGACATTCGG |
| Mouse *Olfr873* target (KD) | GAAATAACCCGAAGAGAAT |
| Mouse *Iars2* target (KD) | GCAGCCGCCCGGTACCTAA |
| Mouse *Olfr524* target (KD) | GAACAGCCCTAGAAGGCTT |
| Mouse *Sod1* target (KD) | GCCTCCCCGCGCCCCGGAG |
| Mouse *Got1* target (KD) | TGTACACCGGAGGAGGTGT |
| Mouse *Syngr2* target (KD) | CGAGGCGCGTCACCACCTG |
| Mouse *Extl2* target (KD) | GAGGGAGGGGTGGCCACTC |
| Mouse *Atp13a2* target (KD) | TCCGCGGAGGGGCCGGGCT |
| Mouse *Zyg11b* target (KD) | GCTCCAGGAGGCTCGGGTC |
| Mouse *Arl8b* target (KD) | GGGCCGATACCCGCTCGT |
| Mouse *Hgs* target # 1 (KO) | TCTGCGACCTGATCCGTCAG |
| Mouse *Hgs* target # 2 (KO) | TCCGGAACGAACCCAAGTAC |
| Human *Hgs* target # 1 (KO) | GTACACTTCGTACCCCAAGG |
| Human *Hgs* target # 2 (KO) | ACTCTTCATGCGGTTCACGA |
| Monkey *Hgs* target # 1 (KO) | CTTGGGGTACGAAGTGTACG |
| Monkey *Hgs* target # 2 (KO) | CCGGAATGAGCCTAAGTACA |

**Table S6.** List of antibodies.

| Antibodies |  |  |
| --- | --- | --- |
| APC anti-human CD107a (LAMP-1) Antibody | BioLegend | Cat# 328620  (FC:1/100) |
| FITC anti-human CD107a (LAMP-1) Antibody | BioLegend | Cat# 328606  (FC:1/100) |
| CoraLite® Plus 488 Anti-Mouse CD107a / LAMP1 (1D4B) | Proteintech | (IF:1/100) |
| β-actin Mouse Monoclonal Antibody | Beijing Ray Antibody Biotech | Cat# RM2001-IR  (WB:1/5000) |
| HRS (D7T5N) Rabbit mAb (Hgs) | Cell Signaling Technology | Cat# 15087S  (WB:1/2000; IF:1/500; IP:1/250) |
| CoraLite® Plus 488-conjugated HGS Polyclonal antibody | Proteintech | Cat# CL488-10390  (IF:1/100) |
| ERGIC53 Rabbit pAb A10440 | Abclonal | Cat# A10440  (IF:1/100) |
| GOLGA2/GM130 Monoclonal antibody | Proteintech | Cat# 66662-1-Ig-100ul  (IF:1/100) |
| β-tubulin Mouse Monoclonal Antibody | HUABIO | Cat# EM0103  (WB:1/5000) |
| GST Tag Monoclonal antibody | Proteintech | Cat# 66001-2-1g  (WB:1/2000) |
| Anti-HA-tag mAb | MBL | Cat# M180-3  (WB:1/2000; IF:1/250) |
| DYKDDDDK Tag Monoclonal Antibody | Proteintech | Cat# 66008-4-1g  (WB:1/2000; IF:1/250) |
| GFP-tag Mouse Monoclonal Antibody | Ray Antibody | Cat# RM1008  (WB:1/5000) |
| SARS-CoV / SARS-CoV-2 (COVID-19) spike antibody [1A9] | Genetex | Cat# GTX632604-S  (WB:1/2000) |
| SARS-CoV-2 S protein (944-1214 aa) Polyclonal antibody | Proteintech | Cat# 28867-1-AP  (IF:1/100) |
| SARS-CoV-2 (COVID-19) Membrane antibody [HL1088] | Genetex | Cat# GTX636246-S  (WB:1/2000; IF:1/250) |
| SARS-CoV/SARS-CoV-2 Nucleocapsid Antibody, Rabbit MAb | Sino Biological | Cat# 40143-R004  (WB:1/2000) |
| SARS-CoV-2 (COVID-19) Nucleocapsid antibody, Human MAb | This paper | (IF:1/100) |
| Human coronavirus (HCoV-HKU1) Nucleocapsid Antibody, Rabbit PAb | Sino Biological | Cat# 40642-T62  (WB:1/1000; IF:1/100) |
| Human coronavirus (HCoV-OC43) Nucleocapsid Antibody, Rabbit PAb | Sino Biological | Cat# 40643-T62  (WB:1/1000; IF:1/100) |
| Human coronavirus (HCoV-229E) Nucleocapsid Antibody, Rabbit PAb | Sino Biological | Cat# 40640-T62  (WB:1/1000; IF:1/100) |
| Human coronavirus (HCoV-NL63) Nucleocapsid Antibody, Rabbit PAb | Sino Biological | Cat# 40641-T62  (WB:1/1000; IF:1/100) |
| MERS-CoV Nucleocapsid Protein Antibody, Rabbit PAb | Sino Biological | Cat# 40068-RP02  (WB:1/1000) |
| SARS-CoV-2 (COVID-19) Envelope antibody [HL1443] | Genetex | Cat# GTX636915-S  (WB:1/1000) |
| MHV Nucleocapsid Antibody, Mouse MAb [2E6] | Gifted by Rong Ye | N/A |
| Alexa Fluor 488 Anti-Cytokeratin 5 antibody [EP1601Y] | Abcam | Cat# ab193894  (IF:1/400) |
| Anti-alpha Tubulin (acetyl K40) antibody [6-11B-1] | Abcam | Cat# ab24610 |
| MUC5AC Monoclonal Antibody (45M1) | Thermo Fisher | Cat# AC011 |
| Mouse Control IgG | Abclonal | Cat# AC011 |
| Rabbit Control IgG | Abclonal | Cat# AC005 |
| Goat anti-Mouse IgG (H+L) Highly Cross-Adsorbed Secondary Antibody, Alexa Fluor Plus 488 | Invitrogen | Cat# A32723 |
| Goat anti-Rabbit IgG (H+L) Highly Cross-Adsorbed Secondary Antibody, Alexa Fluor Plus 488 | Invitrogen | Cat# A32731 |
| Goat anti-Mouse IgG (H+L) Cross-Adsorbed Secondary Antibody, Alexa Fluor 568 | Invitrogen | Cat# A11004 |
| Goat anti-Rabbit IgG (H+L) Cross-Adsorbed Secondary Antibody, Alexa Fluor 568 | Invitrogen | Cat# A11011 |
| Goat anti-Mouse IgG (H+L) Highly Cross-Adsorbed Secondary Antibody, Alexa Fluor Plus 647 | Invitrogen | Cat# A32728 |
| Goat anti-Rabbit IgG (H+L) Highly Cross-Adsorbed Secondary Antibody, Alexa Fluor Plus 647 | Invitrogen | Cat# A32733 |
| Goat anti-Human IgG (H+L) Cross-Adsorbed Secondary Antibody, Alexa Fluor 488 | Invitrogen | Cat# A11013 |
| IRDye® 800CW Goat anti-Rabbit IgG Secondary Antibody | LI-COR | Cat# 926-32211 |
| IRDye® 680RD Goat anti-Mouse IgG (H+L) | LI-COR | Cat# 926-68070 |

**Table S7.** List of Critical commercial assasy.

| EasyPure® Simple Viral DNA/RNA Kit | TransGen Biotech | Cat# ER211-01 |
| --- | --- | --- |
| NucleoBond Xtra Midi (50) | MACHEREY-NAGEL | Cat# 740410.50 |
| ClonExpress® Ultra One Step Cloning Kit | Vazyme Biotech | Cat# C115-01 |
| Phanta® Max Super-Fidelity DNA Polymerase | Vazyme Biotech | Cat# P505-d2 |
| SuperScript™ IV First-Strand Synthesis System | Invitrogen&trade | Cat# 18091050 |
| Cell Counting Kit-8 | Beyotime | Cat# C0039 |
| BD Cytofix/Cytoperm™ solution | BD Bioscience | Cat# 554722 |
| 1 × BD Perm/Wash buffer | BD Bioscience | Cat# 554723 |
| NucleoSpin® Blood | MACHEREY-NAGEL | Cat# 740951.50 |
